# Unveiling causal regulatory mechanisms through cell-state parallax

**DOI:** 10.1101/2023.03.02.530529

**Authors:** Alexander Po-Yen Wu, Rohit Singh, Christopher Walsh, Bonnie Berger

## Abstract

Genome-wide association studies (GWAS) identify numerous disease-linked genetic variants at noncoding genomic loci, yet therapeutic progress is hampered by the challenge of deciphering the regulatory roles of these loci in tissue-specific contexts. Single-cell multimodal assays that simultaneously profile chromatin accessibility and gene expression could predict tissue-specific causal links between noncoding loci and the genes they affect. However, current computational strategies either neglect the causal relationship between chromatin accessibility and transcription or lack variant-level precision, aggregating data across genomic ranges due to data sparsity. To address this, we introduce GrID-Net, a graph neural network approach that generalizes Granger causal inference to detect new causal locus–gene associations in graph-structured systems such as single-cell trajectories. Inspired by the principles of optical parallax, which reveals object depth from static snapshots, we hypothesized that causal mechanisms could be inferred from static single-cell snapshots by exploiting the time lag between epigenetic and transcriptional cell states, a concept we term “cell-state parallax.” Applying GrID-Net to schizophrenia (SCZ) genetic variants, we increased variant coverage by 36% and uncovered noncoding mechanisms that dysregulate 132 genes, including key potassium transporters such as KCNG2 and SLC12A6. Furthermore, we discovered evidence for the prominent role of neural transcription-factor binding disruptions in SCZ etiology. Our work not only provides a strategy for elucidating the tissue-specific impact of noncoding variants but also underscores the breakthrough potential of cell-state parallax in single-cell multiomics for discovering tissue-specific gene regulatory mechanisms.

## Introduction

Dynamic changes in the chromatin states of cis-regulatory genomic elements influence gene expression in many biological processes, including development, differentiation, reprogramming, and perturbation response^1–3^. Correspondingly, genome-wide association studies (GWAS) have implicated thousands of noncoding loci in diseases and complex traits^4,5^. Yet, the challenge of linking these noncoding loci to the genes they regulate has impeded therapeutic advances. The common approach of simply associating a locus to its nearest gene is inadequate, as it overlooks distal locus-gene interactions. This is a critical limitation, as distal interactions are likely to be important for many genes. Meanwhile, experimental approaches to discover such interactions can be difficult to scale. Expression quantitative trait locus (eQTL) analyses require sample collections from large cohorts^6^, and CRISPR perturbations have been limited to specific loci^7^. Computational methods based on bulk data have also been proposed, such as Wang et al.’s integration of multiple data types to link variants to genes^8^, and Fulco et al.’s activity-by-contact (ABC) model^9,10^ that predicts enhancers of a gene based on the activity and 3D chromatin structure around the gene. However, these models focus on the relative spatial positions of enhancers and genes, and are not designed to evaluate whether changes in accessibility at a locus actually influence gene expression.

Recent advances in single-cell multimodal technology, which can simultaneously profile chromatin accessibility (scATAC-seq) and gene expression (scRNA-seq) in the same cell, offer a promising opportunity to uncover tissue-specific, genome-scale, causal relationships between noncoding loci and genes in a single experiment^11–14^. However, current computational methods face substantial hurdles in extracting meaningful locus-gene associations. Existing approaches have diverse and often unrelated objectives: achieving concordance between the modalities^15,16^, inferring gene regulatory networks by considering the accessibility of transcription-factor binding motifs^17,18^, or using chromatin accessibility to improve RNA velocity inference^19^. Unfortunately, owing to the extreme sparsity of scATAC-seq data (< 1% non-zero entries per cell ^20^), these approaches aggregate chromatin peaks within a defined range of a gene; for instance, MultiVelo consolidates chromatin signals into a single, per-gene score^19^. This results in a loss of variant- and locus-level resolution and also risks ignoring distal peaks. Moreover, these methods are largely reliant on correlational analyses between scATAC-seq and scRNA-seq, thereby overlooking the inherent causal time lag in gene regulation (**Figure 1a**). For instance, poised enhancers, which play essential roles in development, undergo changes in chromatin states and histone modifications *prior* to activating their target genes and would be missed by a correlational analysis^21^. Consequently, the appropriate epigenetic and transcriptional readouts to be cross-referenced may correspond to subtly different cell states.

**Figure 1.**
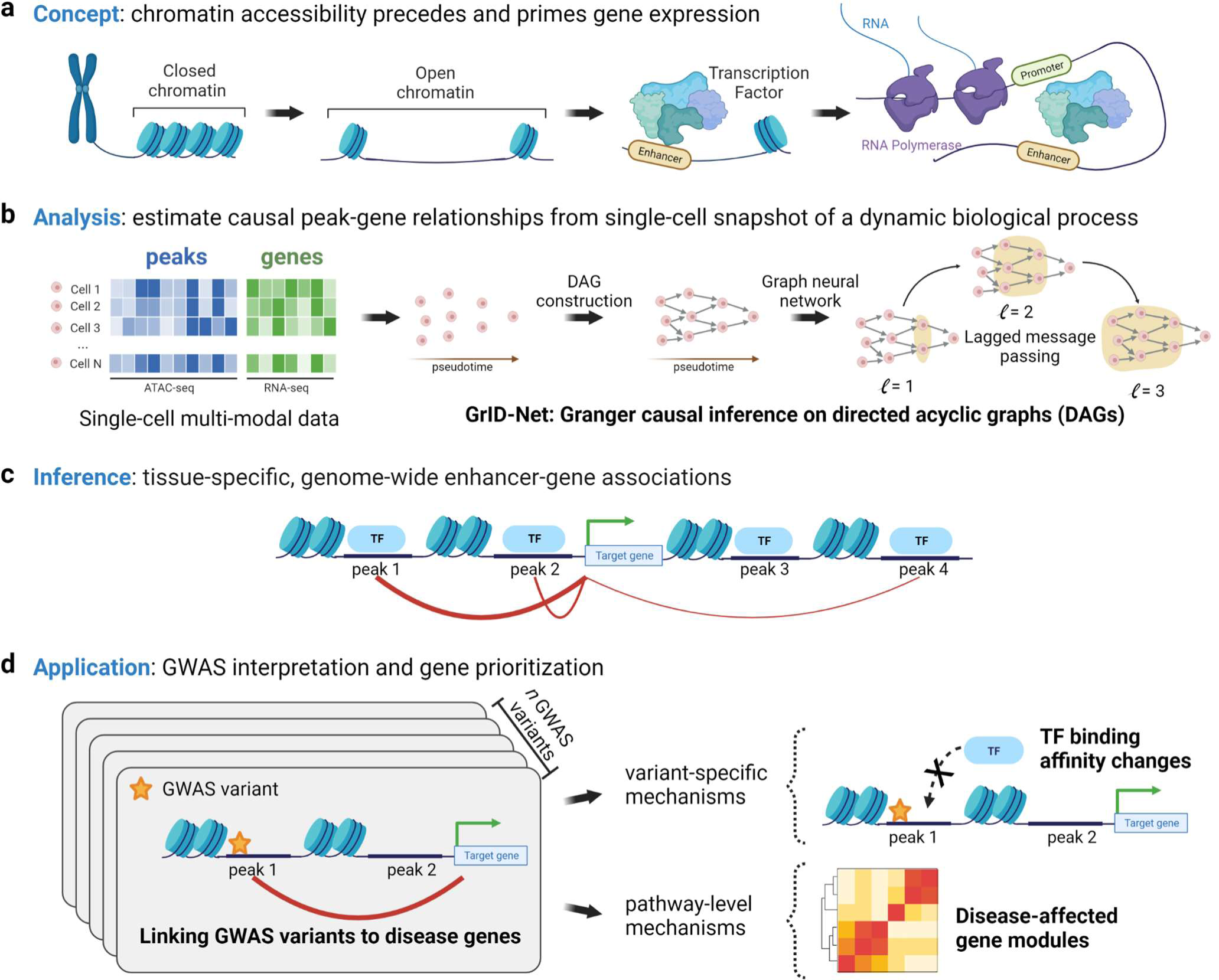
GrID-Net Framework. **(a)** The causal asynchrony between chromatin accessibility and gene expression results in “cell-state parallax”, where a temporal lag exists between these two modalities. Leveraging this parallax from single-cell snapshots enables inference of causal locus-gene associations. **(b)** To identify links between noncoding loci (i.e. peaks) and the genes they regulate, we apply GrID-Net to single-cell multimodal data in which chromatin accessibility (ATAC-seq) and gene expression (RNA-seq) have been co-assayed in the same cell. GrID-Net is an extension of classical Granger causality that can operate on DAG-based representations of single-cell trajectories. The core of GrID-Net is a graph neural network-based lagged message passing architecture. **(c)** GrID-Net enables the detection of asynchronous interactions between regulatory activity at accessible noncoding loci and their downstream effects on gene expression. **(d)** GrID-Net peak-gene links enable the functional interpretation of genetic variants associated with disease. We present a framework for studying the functional effects of such variants both at the level of specific variants and with respect to disease-affected modules.

We refer to this temporal discrepancy as “cell-state parallax,” drawing inspiration from the optical concept of parallax, which describes how different vantage points can unveil new insights. Cell phone cameras with dual lenses leverage optical parallax by capturing multiple 2D images of the same object from slightly offset positions to enable 3D depth perception. Analogously, in single-cell multimodal assays, each modality captures a snapshot of the cell’s state, but at an offset due to the temporal lag between the epigenome and the transcriptome. By exploiting this offset, causal relationships between the modalities can be unveiled. However, a fundamental challenge in accounting for these offsets is that single-cell experiments destroy cells upon measurement, so the temporal ordering of single-cell measurements is often unclear. Moreover, an inference framework needs to be robust to the sparsity of both scATAC-seq and scRNA-seq data^22,23^. Some previous methods mitigate sparsity by averaging transcriptionally similar cells into pseudocells^11^, but this comes at the cost of losing cell-level granularity which is crucial for discerning subtle offsets between distinct modalities. In this work, we operate at a per-peak and per-cell resolution, addressing sparsity through our machine learning framework.

To resolve single-cell multimodal snapshots into causal regulatory relationships, we leverage mathematical principles that underlie the dynamics between causes and their effects. We focus on Granger causality, a statistical framework which exploits the idea that a cause precedes, and is therefore predictive of, its downstream effect. There, a temporally causal relationship between two time-varying processes can be inferred solely by autoregressive analysis of time-resolved observational data. Granger causality has been a powerful tool in biology, econometrics, and other domains for causal discovery without the need to perform interventions.

However, the classical framework of Granger causality is limited to settings where observations can be ordered along a single time-axis. Mathematically, an ordering of observations can be categorized as *total* or *partial*: in the former, any pair of observations can be ordered unambiguously while for the latter only a subset of observations can be so ordered. Partial orderings characterize many dynamical systems, including disinformation cascades in social media, financial transaction networks, and parallel computing systems. In single-cell biology, when a differentiation trajectory has multiple branches, cell states across distinct branches are not directly comparable, necessitating a partial ordering of cells. Classical Granger causal analysis, being limited to total orderings, therefore needs to be adapted for inferring causal single-cell dynamics.

Our key mathematical advance is a generalization of Granger causality to partial orderings. We exploit the equivalence between such orderings and directed acyclic graphs (DAGs) and design a graph neural network framework to infer causality from the dynamics. We complement this with the conceptual advance of cell-state parallax from which we infer a DAG-based representation of single-cell multimodal dynamics. We combine these innovations into GrID-Net (Granger Inference on DAGs), a novel framework for Granger causal inference on DAGs that we apply to single-cell multimodal data for inferring causal noncoding locus-gene relationships (**Figure 1b,c**). Leveraging lagged message-passing on a graph neural network, GrID-Net resolves the cell-state parallax between the epigenome and transcriptome. We demonstrate that it outperforms existing state-of-the-art approaches for accurately identifying noncoding locus-gene links across a variety of cell types and single-cell multimodal assays. We then use GrID-Net to link fine-mapped genetic variants associated with schizophrenia (SCZ) to the genes they affect. We uncover noncoding mechanisms of gene dysregulation that prominently feature disruptions of neural transcription-factor binding and synaptic pathways as putative drivers of SCZ disease pathogenesis (**Figure 1d**). Our findings demonstrate the power of GrID-Net in uncovering tissue-specific causal mechanisms of disease and development, using data generated from just one sample. Our work also illustrates the utility of multimodal single-cell studies for investigating the impact of genetic variants on function and phenotype.

The code for GrID-Net can be found at https://github.com/alexw16/gridnet and the full list of SCZ variant-gene links can be found in **Supplemental Table 1**.

### Overview of GrID-Net

In general, inferring true causality can be a challenge as it requires perturbations to confirm cause-and-effect relationships and validations to rule out potential confounders. Granger causality is a powerful statistical concept that can *approximate* causality in time-varying processes solely from observational data. It leans on the principle that a cause precedes its effect^24,25^. Given two time-dependent variables *x* (a potential cause) and *y* (the effect), if the forecast for *y* can be improved by accounting for the past values of *x*, then *x* is said to Granger cause *y* since changes in *x* temporally precede those in *y*.

One of our key contributions is to broaden the set of domains to which Granger causal analyses can be applied. Specifically, we generalize Granger causal inference to dynamical systems that are characterized by partial orderings. To model the asynchrony between cause and effect, we modify standard graph neural networks to allow for lagged message passing. Each node (i.e., a cell state) in the directed acyclic graph accumulates information from its parent nodes (past states), leveraging this lagged information to forecast the present state in the context of the graph-structured dynamics. GrID-Net consists of two forecasting models: a *reduced* model forecasts an effect *y* using only the past of *y* and a *full* model forecasts the effect *y* using both the past of *y* and the past of the candidate causal variable *x*. The strength of the Granger causal link between *x* and *y* can be evaluated by the statistical F-test which quantifies the improvement in forecasting accuracy of the *full* model over that of the *reduced* model.

GrID-Net enables the detection of asynchronous interactions between gene regulatory elements along single-cell trajectories. For each single-cell multimodal dataset, a DAG of cell states is constructed from a k-nearest neighbor graph of cells, with edges oriented in accordance with pseudotime computed after a multimodal integration with Schema^20^. RNA-velocity based edge orientations that capture local dynamics can also be supplied, so long as any resulting cycles are broken. We evaluated the Granger causal strength between each gene and chromatin accessible element (“peak”) within one megabase (Mb) of the gene’s transcription start site, ranking the peak-gene pairs by their Granger causal strengths (**Methods**).

### GrID-Net more accurately identifies noncoding locus-gene links

We evaluated GrID-Net on four single-cell multimodal datasets (scATAC-seq and scRNA-seq) that characterize a range of biological processes, including cell differentiation and drug treatment response^11,12,26,27^. For each dataset, we sought to validate against expression quantitative trait (eQTL) loci profiled in matching tissue (**Methods**). As eQTLs represent associations between genetic variation at a locus and the corresponding changes in a gene’s expression, they serve as useful genome-wide proxies for perturbational validation of causal interactions between genomic loci and genes.

For each gene, we intersected all ATAC-seq peaks within 1 Mb of its transcription start site (TSS) with the set of eQTL variants linked to it, associating each peak with the most significant variant overlapping it. We then evaluated the peak–gene pairs prioritized by GrID-Net and two correlation-based methods used in previous work^11,27^, Pearson correlation and pseudocell correlation. The latter computes correlations on measurements aggregated across multiple cells. We also evaluated the ABC model’s predictions on bulk data from matching tissues, comparing it against the predictions from the single-cell datasets (**Methods**).

We evaluated the accuracy of each method in retrieving the peak–gene pairs that overlap significant eQTLs (*P* < 10^−10^); our findings were robust to the choice of the eQTL significance threshold (**Figure S1**). GrID-Net was significantly more accurate than the other methods across all datasets (**Figure 2a**), exhibiting 7-27% higher area under the receiver operating characteristic curve (AUROC) in 3 of the 4 datasets. GrID-Net demonstrated the smallest improvement in AUROC for the sci-CAR dataset, although all methods performed poorly on this dataset, likely due to the lower data quality of sci-CAR as the earliest single-cell multimodal assay developed. Meanwhile, the correlation-based methods exhibited near-random performance across all datasets, consistently underperforming not only GrID-Net but also the bulk data-based ABC model. The low accuracy of predictions by correlation-based methods, contrasted with the substantial improvement offered by GrID-Net, demonstrates the potential to unravel much richer regulatory insights from these multimodal assays than has been possible.

**Figure 2.**
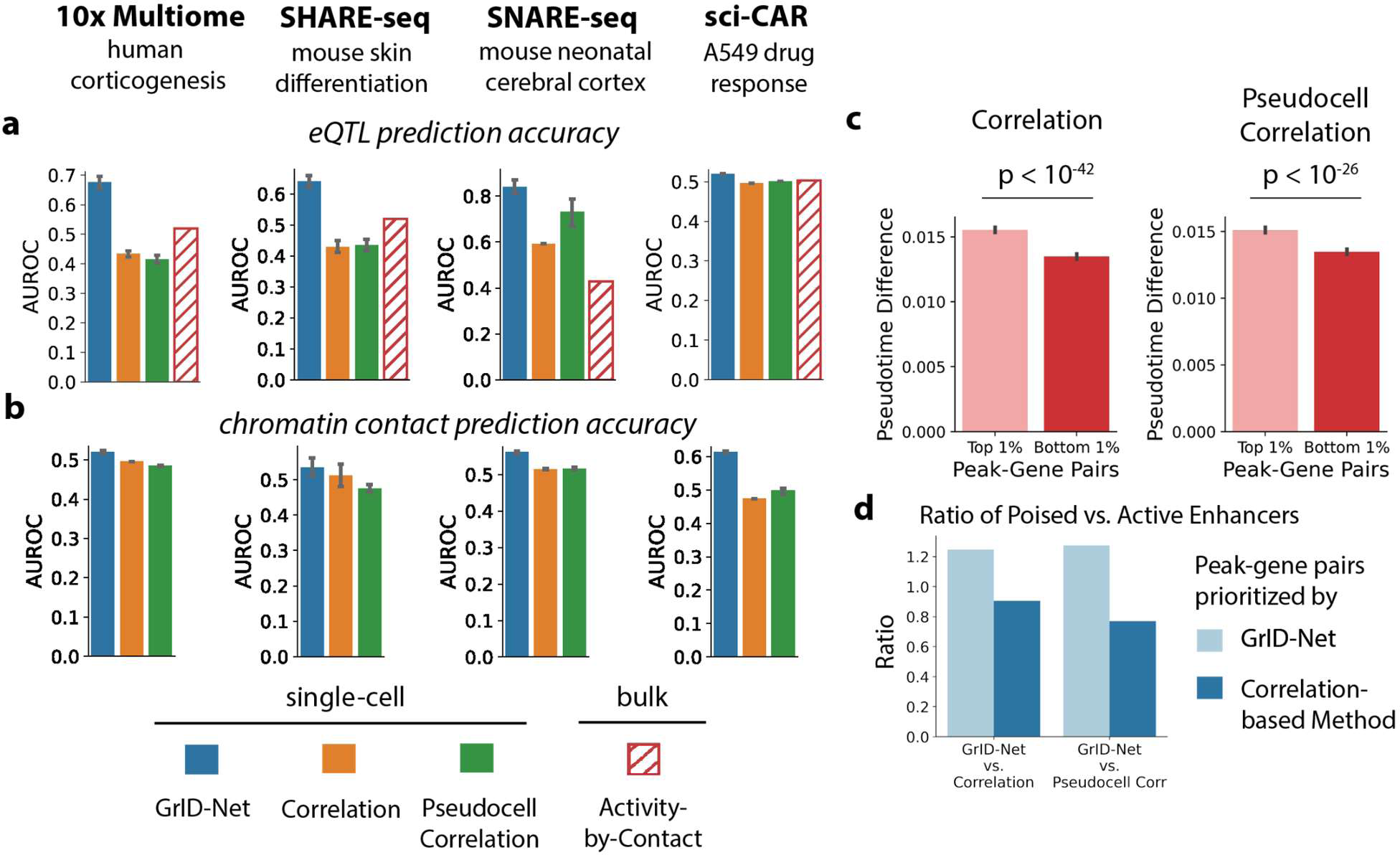
GrID-Net links noncoding loci to genes from single-cell multimodal dynamics. **(a)** Accuracy of single-cell (GrID-Net, Pearson correlation, and pseudocell correlation) and bulk (Activity-By-Contact) methods in predicting expression quantitative trait loci (eQTLs) in matching tissue. In this evaluation, we use eQTLs as proxies for causal interactions between genomic loci and genes. **(b)** Accuracy of methods in predicting high frequency three-dimensional chromatin contacts. The Activity-By-Contact (ABC) model is excluded here since it uses chromatin interaction data as one of its inputs. **(c)** Compared to existing approaches, GrID-Net is better able to identify asynchronous peak-gene associations. On the human corticogenesis dataset, we collected peak-gene pairs that GrID-Net prioritized differently than the correlation-based approaches (top 1%: preferred by GrID-Net; bottom 1%: preferred by the other approach). For each pair, we quantified the asynchrony as the separation between pseudotime values corresponding to the cells where the peak and gene achieved their maximum values, respectively. **(d)** GrID-Net effectively captures instances of chromatin priming. On the human corticogenesis dataset, we show the ratio of poised enhancers to active enhancers for peaks in peak-gene pairs prioritized by GrID-Net vs. each of the correlation-based methods. The estimation of poised-vs.-active enhancers was done using publicly-available histone modification data.

We next assessed the robustness of the single-cell methods to sparsity in single-cell data. GrID-Net better handled sparse data (**Figure S2**), yielding higher accuracies than the correlation-based methods for peak–gene predictions involving genes with sparse data (non-zero transcripts in fewer than 10% of cells) or sparse peaks (under 2% of cells). Notably, pseudocell correlation failed to consistently outperform standard Pearson correlation on these sparse peak–gene cases, even though the former is explicitly designed to compensate for the sparsity of measurements at the level of individual cells. This may be because pseudocell averaging reduces the effective number of independent observations and thus relinquishes the statistical advantages of single-cell heterogeneity. Altogether, these analyses indicate not only GrID-Net’s robustness to the current sparsity of single-cell data but also the potential for its increased effectiveness as the sensitivity of these multimodal assays continues to improve.

To further benchmark GrID-Net using functional indicators of noncoding locus–gene interactions, we examined three-dimensional chromatin contacts (from bulk Hi-C data) between gene promoters and noncoding loci. Compared to the two correlation-based methods, we found that GrID-Net more accurately prioritized peak–gene pairs that were in frequent three-dimensional contact (**Figure 2b)**. The ABC method uses bulk Hi-C data itself as an input so we did not evaluate its agreement with chromatin contacts. In future, the GrID-Net framework could be expanded to also utilize Hi-C data when available— as we describe later, incorporating one-dimensional sequence proximity as a signal into GrID-Net improves its results. However, Hi-C data is currently available only for a limited set of tissues^28^ and the ability to make regulatory inferences from just one multimodal assay enables the convenient study of rare cell types, cancer mutations and drug responses etc.

### GrID-Net reveals how chromatin accessibility primes gene activity

To better understand the biological basis of GrID-Net’s superior accuracy, we applied it on a single-cell multimodal dataset profiling human corticogenesis. We then honed in on peak–gene pairs where GrID-Net and correlation-based methods disagree. Specifically, peak–gene pairs were ranked according to the difference between their GrID-Net and correlation-based prioritization scores (**Methods**), with higher ranking pairs being the ones that were more strongly prioritized by GrID-Net. We then investigated the distinguishing biological characteristics of these differentially-prioritized pairs.

One of our motivations behind GrID-Net is the observation that regions of chromatin around lineage-determining genes become accessible prior to gene expression, foreshadowing lineage choice^26,29^. GrID-Net was designed to resolve this cell-state parallax between chromatin accessibility and gene expression and we assessed if it was able to do so. We first associated each peak and each gene with the pseudotime corresponding to its highest value to determine the lag associated with each peak-gene pair. For both correlation-based methods, we found that peak–gene pairs differentially prioritized by GrID-Net are marked by significantly longer pseudotime lags between chromatin accessibility and gene expression (*P* < 10^−26^, Welch’s one-tailed *t*-test, **Figure 2c**, **Methods**). This result supports GrID-Net’s ability to account for the asynchrony between noncoding regulatory activity and downstream gene expression control that are expected to characterize causal interactions between the two. To explore the regulatory implications of this asynchrony, we obtained data on histone modification states of enhancers in neural progenitor cells and cross-referenced it with ATAC-seq peaks^30^. In peak–gene pairs differentially prioritized by GrID-Net, more poised enhancers (H3K4me1 only) overlapped with the peaks than active enhancers (both H3K4me1 and H3K27ac)^31^ (**Figure 2d**, **Methods**). This was not the case for peak–gene pairs differentially prioritized by correlation-based methods. Poised enhancers are regulatory sites that can trigger future gene expression changes but have not yet affected expression^31,32^; in contrast, active enhancers correspond to ongoing transcription^33^. This suggests that resolving cell-state parallax enables the detection of lagged relationships between the activities of enhancers and their target genes. GrID-Net can thus leverage the concept of chromatin regulatory potential^26^ to facilitate a deeper understanding of tissue-specific causal regulatory mechanisms.

To evaluate the biological significance of capturing these putative causal interactions, we examined the functions of genes implicated in peak–gene pairs differentially prioritized by GrID-Net. We found these genes to be highly enriched for gene sets associated with neurogenesis regulation and neural progenitor cells (*FDR* < 10^−7^, GSEA Preranked^34^), directly relating to the process of corticogenesis profiled in this dataset. We emphasize that the enrichment score is calculated only on the peak–gene pairs that GrID-Net prioritizes over the correlation-based methods. In comparison, the set of genes from peak–gene pairs that *all* methods score highly is functionally enriched for housekeeping genes rather than neuronal activity, suggesting that GrID-Net’s ability to detect poised enhancers is key to discovering cell-type specific regulatory relationships. Gene expression control during differentiation has been reported to depend highly on asynchronous enhancer-gene interactions^35^. The enrichment of these differentiation-related gene sets thus underscores GrID-Net’s ability to construct accurate genome-wide maps of noncoding locus–gene links that go beyond correlative analyses and towards causality.

### Linking variants to schizophrenia disease genes

The ability to link noncoding loci to the genes they regulate enables the discovery of molecular mechanisms underlying complex diseases. Genome-wide association studies (GWAS) have identified thousands of loci associated with diseases^36^. However, over 90% of GWAS variants lie in noncoding regions^37^, and interpreting the functions of these noncoding variants involves the challenge of linking them to the specific genes that they regulate in a cell type-specific manner. With GrID-Net, we can leverage data from a single-cell assay on matching tissue to uncover these cell-type-specific variant-gene links and help bridge the gap between genetic variants and gene dysregulation in disease.

We used GrID-Net to study variants associated with schizophrenia (SCZ), a psychiatric disorder for which inherited genetic variants are believed to underlie its pathogenesis^38^. SCZ’s high prevalence of around 1%, its substantial morbidity and mortality, and the lack of effective treatments are largely due to our limited understanding of SCZ pathophysiology^39,40^. We applied the GrID-Net peak-gene links inferred from human corticogenesis^27^, to interpret and analyze variants reported in the largest GWAS of SCZ to date^41^. We refer to this original study as the Trubetskoy, Pardiñas *et al*. study.

To boost GrID-Net’s accuracy on shorter-range peak–gene interactions, we incorporated one-dimensional genomic distance as an additional feature for predicting peak–gene associations. We observed a higher likelihood of peak–gene interaction if the two are located close to each other, in line with reported patterns of genomic distance-dependent regulatory effects^42,43^. We calibrated the relative importance of this feature on a dataset profiling mouse skin differentiation^26^ where we fitted a logistic regression model that combines the raw GrID-Net score with a genomic distance score (**Methods**). We found that integrating genomic distance with GrID-Net improved predictions of peak–gene links for mouse skin cells. The discriminative power on short-range peak–gene interactions improved, while the ability to detect long-range interactions was preserved. Furthermore, this model trained on mouse skin cell data performed well across species and cell types in human corticogenesis as well (**Figure S3**). This genomic distance-dependent model’s scores were used for all remaining analyses of the human corticogenesis dataset.

From the SCZ GWAS study, we retrieved all fine-mapped genetic variants with posterior probability (*PP*) over 5%. We intersected these variants with peaks from the human corticogenesis dataset. We then generated an initial set of candidate SCZ genes by associating each variant with up to 3 of its overlapping peak’s highest scoring genes, retaining only variant– gene associations with scores in the top 50% of all peak–gene scores. This approach yielded a set of 132 genes that not only is enriched for genes implicated in the original SCZ GWAS study (*FDR* = 0.037, Enrichr^44^) but also includes novel SCZ gene candidates (**Supplemental Table 1**). For example, of the 12 SCZ variants that were prioritized by both GrID-Net and Trubetskoy, Pardiñas *et al*., 9 were linked to the same target genes by both approaches (**Figure 3a)**. This overlap included the association between rs324017 and NAB2, a gene implicated in stress-induced neuroprotective maintenance that also is the downstream target of the SCZ drug cinnarizine^45,46^. For these same 12 SCZ variants, GrID-Net also proposed 20 additional SCZ candidate genes, of which 12 (60%) were found to be specifically expressed in brain tissues and thus potentially relevant in SCZ etiology (**Methods**). Some of these additional candidates include NXPH4, a neurexophilin involved in synapse function and cerebellar motor control^47^, and EPN2, an epsin linked to neural fate commitment^48^. Moreover, of the 78 SCZ variants with *PP* over 50%, only 22 (28%) were linked to genes by the original SCZ GWAS study (**Figure 3b**).

**Figure 3.**
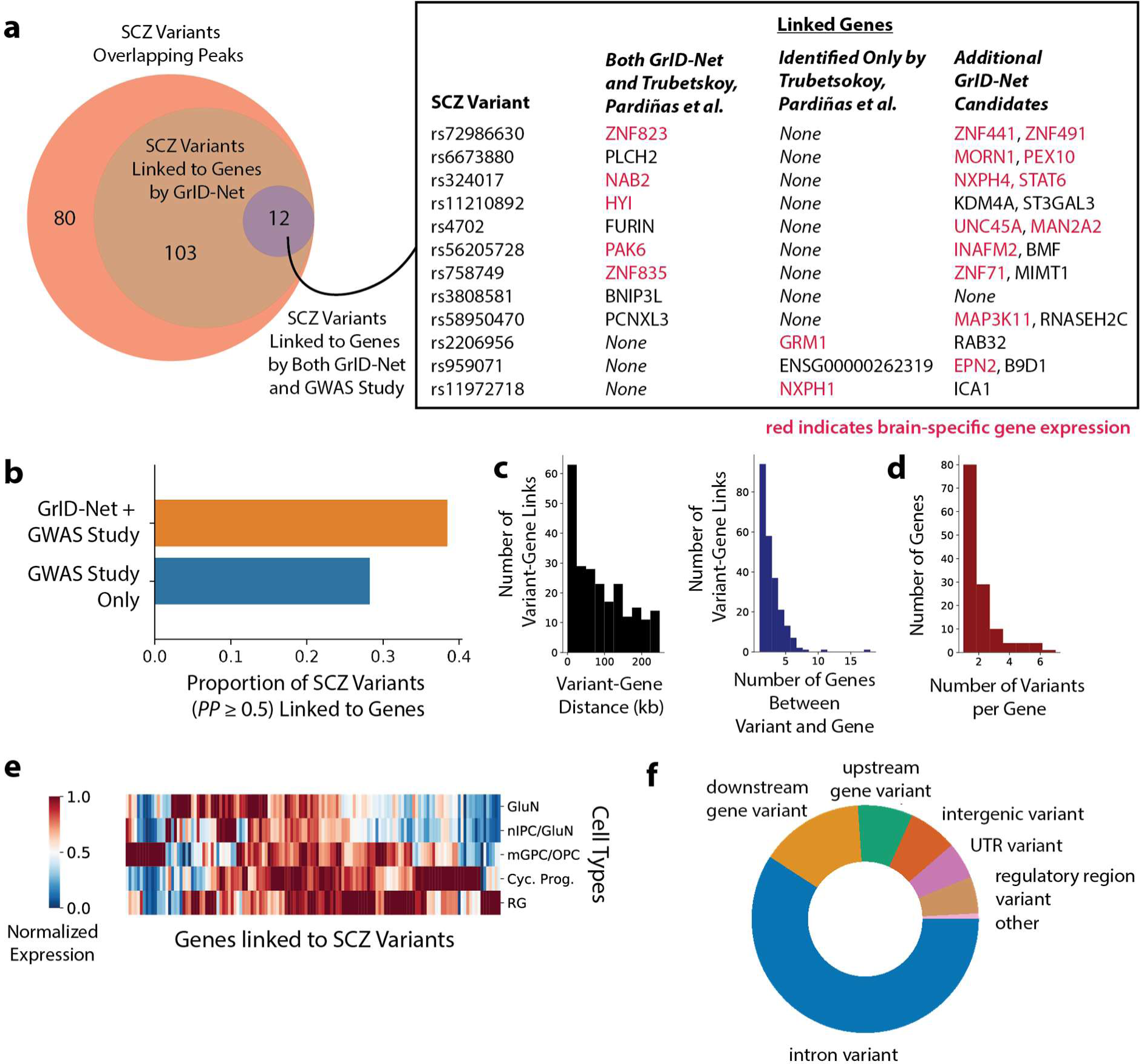
GrID-Net predictions link schizophrenia variants to affected genes. **(a,b)** Schizophrenia (SCZ) variants from Trubetskoy, Pardiñas *et al*., prioritized by GrID-Net. Overall, 195 SCZ variants overlapped with scATAC-seq peaks from the human corticogenesis dataset. Compared to the original study, GrID-Net enables the annotation of many additional variants in this subset. Furthermore, as multimodal single-cell assays become common, we expect GrID-Net’s coverage to widen. We also compared specific SCZ variant-gene links for the 12 variants where the original study also reported an annotation. **(c)** GrID-Net is able to identify long-range variant gene associations: (Left) genomic distance between an SCZ variant and the transcription start site (TSS) of its affected gene(s). (Right) number of genes in the genomic region between an SCZ variant and its affected gene(s). **(d)** Histogram of the number of variants linked to each SCZ gene. **(e)** Genes linked to SCZ variants are cell type-specific. We show the heatmap of normalized expression values for all genes linked to SCZ variants in cell types involved in human corticogenesis. **(f)** Distribution of annotated genomic locations for SCZ variants linked to genes. A substantial fraction of genes identified by GrID-Net are associated with intronic variants which have been reported to have tissue-specific regulatory effects.

Inclusion of SCZ variants that were linked to genes by GrID-Net increased this total to 30 (38%) SCZ variants. GrID-Net can, therefore, be used to complement standard strategies for linking variants to genes, such as those used in the original SCZ GWAS study. It can propose gene associations for variants with no annotations and also augment existing annotations by suggesting additional genes that a variant may be affecting.

In particular, GrID-Net helps address a key challenge in GWAS analyses: the common strategy of assigning GWAS variants to their closest gene^49^ misses many biologically meaningful variant–gene relationships. By leveraging single-cell heterogeneity, GrID-Net can evaluate longer range interactions. Many of the variant–gene links we identify are characterized by large genomic distances between the variant and the transcription start site (TSS) of the linked gene. Moreover, the variants are often linked to genes that are multiple genes over from their closest gene (**Figure 3c**). For example, DCP1B and NCKAP5L, each of which are connected to only a single variant, are separated from their respective variants by 18 and 11 genes (235 and 242 kilobases (kb)), respectively. DCP1B has been implicated in SCZ, hypomania, and memory performance^50,51^, while NCKAP5L, a cytoskeletal gene involved in neuronal activity^52^, has previously been associated with SCZ as well as amyotrophic lateral sclerosis^53^, indicating possible roles for these genes in neuron-specific dysregulation in SCZ.

In addition, 52 candidate genes (39.4% of the overall set) were associated with more than one variant (**Figure 3d**). These genes may be controlled by the coordinated activity of multiple regulatory elements and co-regulators relevant to SCZ^54^. IGSF9B was linked to 7 noncoding SCZ variants – the most of any gene. Coding variants in IGSF9B have been found to be associated with major depression and the negative symptoms of SCZ^55,56^, underscoring its role in regulating affective symptoms in SCZ. Similarly, KCNG2, a potassium channel-encoding gene, was linked to 6 noncoding SCZ variants, and has been proposed as a drug target for SCZ given the known disruption of KCN-related pathways in SCZ^40,57,58^. Neither of these genes were prioritized in the original GWAS analysis.

We also assessed the cell-type specificity of the genes linked to SCZ variants. We found that 104 candidate SCZ genes (78.8% of the overall set) are differentially expressed across the cell types profiled in the human corticogenesis dataset (Welch’s *t*-test with Benjamini-Hochberg correction *P* < 0.05, **Figure 3e**). Further analysis of the SCZ variants also revealed that 59% of the variants lie in intronic regions, while only 7% of variants lie in intergenic regions more than 5 kb from any gene (**Figure 3f**, **Methods**). As enhancers with tissue-specific effects are highly enriched in introns while intergenic enhancers more often regulate housekeeping genes^59^, these findings support the potential cell type-specificity of the regulatory disruptions underlying SCZ pathogenesis.

### Transcription factor binding disruptions are implicated in schizophrenia

Transcription factor (TF) binding at the cis-regulatory elements located in noncoding genomic loci is a key mechanism of gene regulation. We observed that SCZ variants with high *PP* are associated with larger TF binding affinity changes (**Figure S4a**). To investigate the dysregulatory role of SCZ variants prioritized by GrID-Net, we therefore evaluated their effect on TF binding. For each variant associated with a gene by GrID-Net, we calculated the difference in the predicted TF binding affinity between the reference allele and the SCZ risk allele (**Methods**, **Supplemental Table 2**). Of the 115 such SCZ variants, 28 (these were associated with 49 target genes) were predicted to cause a 20-fold or stronger change in at least one TF’s binding affinity (**Figure 4a**). The TFs we identified as having the most widespread binding affinity changes are known to be implicated in SCZ, exemplifying GrID-Net’s ability to deliver de novo mechanistic insight. VEZF1, a vasculature-related TF reported to play a role in SCZ dysregulation^60^, displayed diminished binding affinity most frequently (in 5 variants). In addition, KLF15 and KLF6, two Kruppel-like family (KLF) TFs known to regulate neuronal growth and linked to SCZ^61,41^, were each associated with reduced binding affinity in 4 variants (**Figure S4b**). Meanwhile, the TFs with the most widespread increase in binding affinity – SP1, SP3, and KLF16 – have also been reported to be disrupted in SCZ^62,63^ (**Figure S4c**). More broadly, of the TFs with a 20-fold binding affinity change in at least one variant, 57.9% (=33/57) have cell type-specific expression patterns in corticogenesis (Welch’s *t*-test with Benjamini-Hochberg correction *P* < 0.05). Collectively, these results point to genetic disruption of neuron-specific regulatory elements as potential drivers of SCZ pathogenesis.

**Figure 4.**
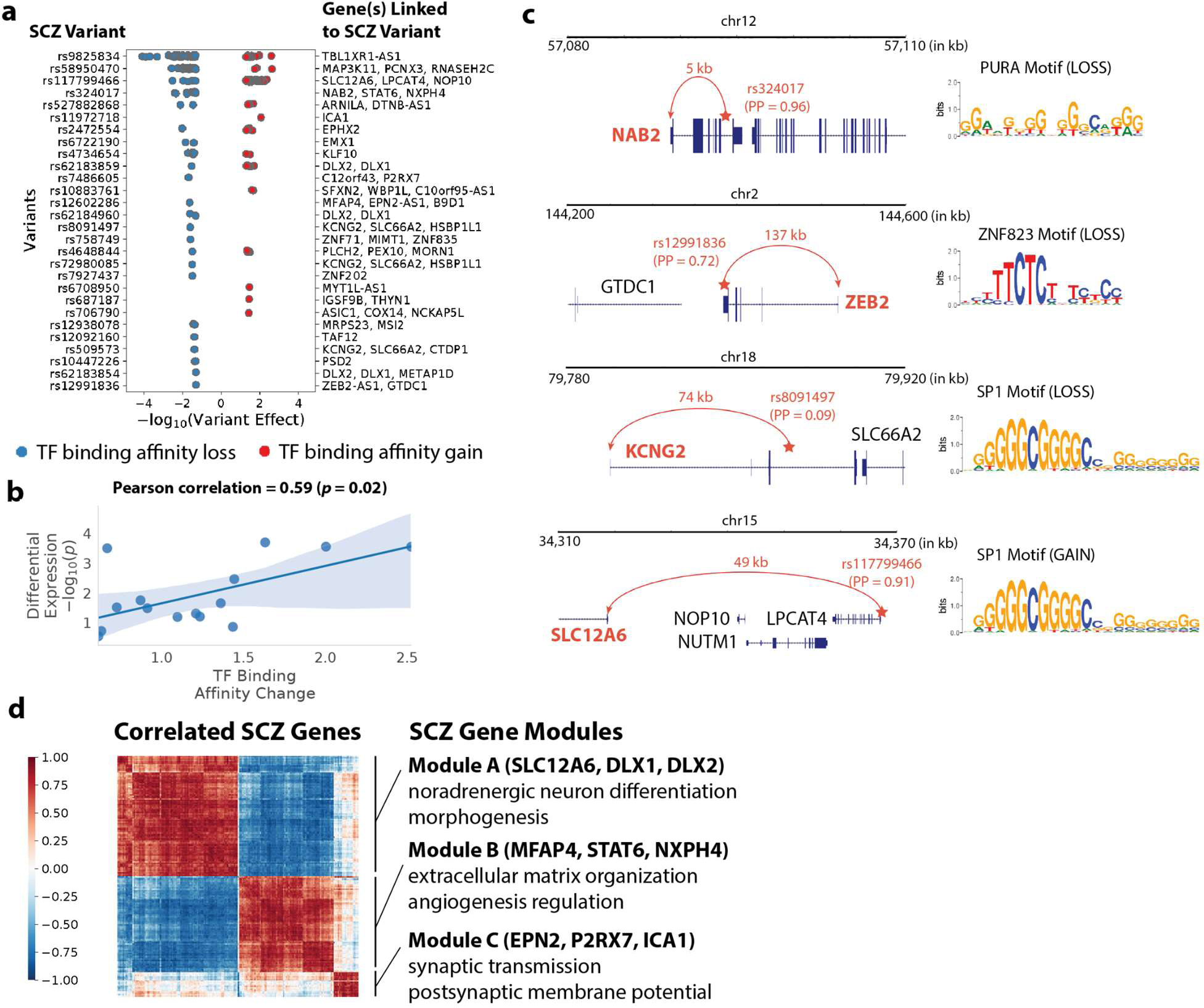
GrID-Net links transcription factor binding disruptions to gene dysregulation in schizophrenia. **(a)** TFs with at least a 20-fold change in predicted binding affinity caused by SCZ variants and the genes linked to these variants. Many of these TFs (e.g., KLF and SP families) are implicated in SCZ. **(b)** Relationship between TF binding affinity changes caused by SCZ variants and significance of differential expression in their linked genes between patient-derived SCZ glial progenitor cells and matching controls. **(c)** Examples of SCZ variant-gene links supported by at least 20-fold changes in predicted TF binding affinity. **(d)** Gene modules co-expressed with SCZ genes in SCZ GPCs and matching controls.

We reasoned that if the TF binding affinity changes at a SCZ variant are phenotypically meaningful, they should result in expression changes of the corresponding gene(s) (as identified by GrID-Net). Accordingly, we obtained gene expression data from glial progenitor cells (GPCs) produced from SCZ patient-derived induced pluripotent cells, as well as matching controls^58,64^, and computed differential expression patterns in SCZ GPCs. Focusing on glial differentiation genes, we observed that the differential expression of a gene in SCZ GPCs was indeed significantly correlated with TF binding affinity changes at the corresponding SCZ variant (Pearson correlation coefficient = 0.59, **Figure 4b**, **Methods**). The consistency between these independent observations provides support to our overall analytic approach and underscores the potential to leverage GrID-Net for examining causal, gene-specific regulatory mechanisms.

We drilled down into specific genes reported by our analysis, focusing on those linked to SCZ variants associated with at least 20-fold TF binding affinity changes (**Figure 4c**). NAB2, the aforementioned gene implicated in neuroprotective processes^46^, was linked to rs324017, which contributes to a 200-fold decline in binding affinity for the TF PURA. Intriguingly, PURA is also critical for neural function and development, and its dysfunction is implicated in several neurological and cognitive impairments^65,66^. In addition, rs12991836 was linked to ZEB2, whose TSS is positioned 137 kb from the variant. ZEB2 has been previously reported as a target of long-range regulation, and heterozygous ZEB2 mutations are known to cause Mowat-Wilson syndrome, which features structural brain abnormalities and intellectual disability^67^.

Furthermore, GrID-Net prioritized two potassium transport genes - KCNG2 and SLC12A6 - a class of genes which has been associated with compromised potassium uptake when dysregulated^58^. This phenotype is characteristic of not only SCZ but also other neuron-related diseases, such as Huntington’s disease, multiple sclerosis, and hemiplegic migraine^64,68^. KCNG2 was linked to 6 SCZ variants, the second most of any gene as described earlier. Using TF binding affinity changes to hone in on the most relevant variants, we identified rs8091497 as the variant with the largest TF binding affinity change. rs8091497 was associated with a 40-fold decline in binding affinity for SP1, which itself has been linked to behavioral abnormalities in SCZ^63^. As for SLC12A6, its corresponding variant (rs117799466) increases the binding affinity for many TFs with repressive capabilities. The three TFs with the greatest binding affinity gains at this locus – SP1, ZN281, and MAZ – all play a role in neuronal differentiation^63,69,70^, and both SP1 and ZN281 have also shown increased expression levels in SCZ as well^71,72^. Combined with their increased binding affinity incurred by rs117799466, the upregulation of these TFs may contribute to the repression of SLC12A6, an outcome that has been previously linked to the potassium uptake impairments that characterize SCZ glia^58^. Together, these findings suggest a possible mechanism for SLC12A6 downregulation in SCZ that implicates enhanced binding of neuron-related repressors to a SLC12A6-targeting enhancer to reduce its expression levels.

We then broadened our analysis to gene modules, exploring if the genes identified by GrID-Net are systematically involved in SCZ-related pathways. The 14 SCZ variants that cause at least 40-fold changes in TF binding affinity are linked by GrID-Net to 26 genes, whose differential expression patterns we examined in SCZ GPCs. We detected 3 distinct gene modules that are co-expressed with several of these genes (**Figure 4d**, **Methods**). Module A (SLC12A6, DLX1, and DLX2) is enriched for genes related to noradrenergic neuron differentiation (*FDR* < 0.05) and morphogenesis (*FDR* < 0.02); Module B (MFAP4, STAT6, NXPH4) is enriched for extracellular matrix organization (*FDR* < 10^−14^) and angiogenesis regulation (*FDR* < 0.02) genes, and Module C (EPN2, P2RX7, ICA1) is enriched for synaptic transmission (*FDR* < 10^−4^) and postsynaptic membrane potential (*FDR* < 0.05) (Enrichr^44^). Each of these pathways has been previously associated with SCZ. Noradrenergic neurons have long been implicated in disrupted neuromodulation in SCZ^73^, neuronal angiogenesis underlies the vascular abnormalities observed in SCZ^74^, and genetic studies have pointed to the critical role of synaptic function in SCZ^41^. These pathways, therefore, enable the interpretation of SCZ variants beyond the dysregulation of the specific genes that they directly affect, presenting an opportunity to discover systems-level mechanisms of SCZ pathogenesis.

## Discussion

Most methods for single-cell multimodal data integration have focused on synchronizing modalities to improve downstream tasks, such as cell type identification. In contrast, we demonstrate it is also powerful to systematically study the asynchrony between the modalities. We presented GrID-Net, a method to uncover causal connections between genes and noncoding genomic loci using co-assayed single-cell RNA-seq and ATAC-seq data. GrID-Net operates at a genome-wide scale, is tissue-specific, and can effectively detect long-range (up to 1 Mb) cis-regulatory associations. A longstanding challenge with GWAS studies has been to identify the specific genes that are dysregulated by noncoding disease-linked variants. With GrID-Net, a single commercially-available assay of the matching tissue can power a genome-wide estimation of variant-gene relationships. By resolving tissue-specific causal variant-gene associations, GrID-Net can help identify disease-linked genes as therapeutic targets.

When studying a particular disease, GrID-Net should be paired with a single-cell assay of a matching tissue undergoing a dynamic biological process, such as development, differentiation, disease, or drug response. GrID-Net can then translate the results of this single assay to genome-wide causal hypotheses specific to the disease. We demonstrate this by leveraging a single-cell assay of human corticogenesis to interpret the functional effects of schizophrenia (SCZ) variants. GrID-Net expands both the breadth and depth of existing gene prioritization analyses: it finds novel associations for many unlinked variants while also uncovering additional causal mechanisms for previously investigated variants. For instance, we identified SCZ variants causing large (>20-fold) changes in TF binding affinity and connected them to 49 genes. Many of these genes are located distally from the SCZ variants that are predicted to affect them and would have been neglected by existing GWAS variant–gene associations analyses that consider only genomic proximity. Furthermore, by correlating TF binding affinity changes with differential expression in SCZ glial progenitor cells, we predicted possible mechanisms of TF-driven dysregulation for genes like KCNG2 and SLC12A6. Combined with an exploration of SCZ-related pathways associated with GrID-Net-identified genes, these analyses provide a comprehensive strategy for unveiling mechanistic insights about SCZ pathogenesis. Such analyses can also be applied to investigate other diseases.

Our work rests on two conceptual insights that are broadly applicable. First, when different modalities are measured in the same cell their causal relationship implies a temporal offset between them, a phenomenon we refer to as “cell-state parallax”. Here, we exploit the parallax between RNA-seq and ATAC-seq, but such parallax can also be leveraged from other co-assayed modalities such as chromatin structure or methylation. We view RNA velocity^75^ and protein velocity^76^ as specific applications of the broader concept of cell-state parallax, the distinction being that velocity uses pre-defined mappings between modalities (e.g., spliced vs. unspliced transcript-counts for a particular gene) to infer the differentiation direction of each cell. In contrast, such mappings between modalities may be unclear in general cell-state parallax and require customized inference approaches. Accordingly, our second conceptual insight is to develop computational frameworks that can accommodate the cross-modality dynamics of cell differentiation. Much of the existing work on modeling gene expression dynamics utilizes mathematical and statistical tools designed for time-series analysis^77,78^, which make the unrealistic assumption that a total ordering of cells can capture the underlying biology. We argue that a partial ordering of cells is necessary to accurately model forks and merges in trajectories and accordingly formulate the differentiation landscape as a directed acyclic graph of cells that can represent such an ordering. We expect a promising direction of future research will be computational inference techniques, like GrID-Net, that can operate under this more realistic model.

Since GrID-Net operates on observational data, the locus-gene associations it infers are causal hypotheses that need to be validated by perturbations. In the eQTL-based evaluation, we sought to mimic such validations by investigating random genetic variants observed in the population and cross-referencing them to observed expression changes. We also note that such causal interactions are likely indirect and mediated by additional biophysical events, such as transcription factor binding. Nonetheless, indirect causality is biologically and therapeutically valuable, as the underlying mechanisms can still be exploited via interventions to achieve the desired phenotype.

With the growing availability of single-cell multimodal datasets, our approach is positioned to serve as a key framework for unlocking new mechanisms of gene regulatory dynamics that underlie fundamental biology and disease. Moreover, GrID-Net can be applied broadly to other domains featuring graph-based dynamics, such as uncovering the drivers of social media disinformation cascades. A key technical advance of this work is generalizing the applicability of standard Granger causal analysis to a wider set of domains than previously possible. Doing so required the use of graph-theoretic deep learning techniques to implement the core statistical intuition of Granger causality for any DAG-structured ordering of observations. Our work, like others before us^79^, thus highlights how classical statistical concepts can be reimagined with modern deep learning techniques to dramatically widen their applicability.

## Methods

### GrID-Net formulation

See Supplementary Note.

### GrID-Net training details and hyperparameters

All GrID-Net models consisted of *L* = 10 GNN layers and were trained using the Adam optimizer with a learning rate of 0.001 and a mini-batch size of 1024 candidate peak-gene pairs. Models were trained for a maximum of 20 epochs or until the relative change in the loss function was less than 0.1/|*P*| between successive epochs. Here, |*P*| denotes the number of candidate peak-gene pairs evaluated per dataset. GrID-Net models were implemented in PyTorch and trained on a single NVIDIA Tesla V100 GPU.

### Single-cell multimodal datasets

We used four single-cell multimodal datasets that spanned a range of different multimodal profiling techniques: 1) a 10x Multiome dataset profiling human corticogenesis, 2) a SHARE-seq dataset profiling mouse skin differentiation, 3) a SNARE-seq dataset profiling mouse neonatal cerebral cortical cells, 4) a sci-CAR dataset profiling A549 cells treated with dexamethasone. For each of these datasets, we followed recommended practice^80^ and applied *log* (1 + *CPM*/100) and *log* (1 + *CPM*/10) transformations to the RNA-seq and ATAC-seq counts data, respectively, where CPM indicates counts per million. We then filtered out all genes that were represented in fewer than 1% of cells in the dataset and all peaks that were represented in fewer than 0.1% of cells. We assembled a set of candidate peak-gene pairs by selecting all peaks with centers within 1 Mb from each gene’s TSS. For the 10x Multiome dataset, we only considered peak-gene pairs that were separated by 250 kb to cohere with the genomic distance threshold used in the original study^27^. To consider multiple samples for each dataset, we generated 5 uniformly sampled sets of 5,000 cells for each of the 10x Multiome, SHARE-seq, and SNARE-seq datasets, while we generated 5 uniformly sampled sets of 3,000 cells for the sci-CAR dataset due its smaller quantity of cells (3,081 cells).

### Directed acyclic graph (DAG) construction

For each sample of a dataset, we first generated per-cell representations for each of the gene expression and chromatin accessibility modalities. Gene expression representations were constructed by (1) scaling the transformed RNA-seq data to unit variance and zero mean, (2) retaining the top 2000 most highly variable genes, (3) and projecting these normalized expression values onto their top 100 principal components (excluding components that have Spearman correlation *ρ* > 0.9 with the total counts per cell. Chromatin accessibility representations were obtained transforming the ATAC-seq data using tf-idf^81,82^ and projecting the resulting values onto the top principal components as in (3) above. We then used Schema^20^ to learn cell representations that unify information from these gene expression and chromatin accessibility representations. We constructed a k-nearest neighbor graph (*k* = 15) based on the Euclidean distances between these cell representations and inferred pseudotime using the scanpy^83^ implementation of diffusion pseudotime^84^ with default parameters. We then removed edges in the k-nearest neighbor graph if they did not align with the direction of increasing pseudotime, thus ensuring that no cycles exist and the resulting graph is a DAG.

### Alternative methods

#### Pearson correlation

We computed the Pearson correlation between the transformed ATAC-seq and RNA-seq values for each candidate peak-gene pair, and the absolute value of the correlation coefficients were used to score the peak-gene pairs.

#### Pseudocell correlation

For each sample, we identified the 49 nearest neighbors for each cell based on the Euclidean distances between the joint representations learned from Schema (see “Directed acyclic graph construction”). We constructed a “pseudocell” from each cell and its 49 nearest neighbors, constructing its RNA-seq and ATAC-seq profiles by summing the raw counts from each of the modalities for all cells included in the pseudocell. We then applied *log* (1 + *CPM*/100) and *log* (1 + *CPM*/10) transformations to the RNA-seq and ATAC-seq counts data, respectively. Pearson correlation was applied to the transformed ATAC-seq and RNA-seq values for each candidate peak-gene pair, and the absolute value of the correlation coefficients were used to score the peak-gene pairs.

#### Activity-by-contact (ABC) model

We obtained ABC prediction scores in cell types matching the cell types profiled in the single-cell multimodal studies. For each single-cell study, we intersected the candidate peak-gene pairs with the ABC enhancer-gene pairs, assigning each peak-gene pair the corresponding ABC score. If no ABC enhancer-gene pair overlap was identified, the peak-gene pair was assigned a value of 0, as the ABC prediction scores were available only for enhancer-gene pairs that exceeded the threshold of 0.015 defined in the original study^10^. For single-cell studies that profiled mouse cells, we mapped ABC enhancer-gene pairs in matching human cell types to the mouse genome due to the strong conservation of epigenomes between human and mouse^85,86^.

### eQTL evaluations

We obtained eQTL data from the eQTL Catalogue^87^ and used eQTL data specific to the cell type profiled in each single-cell multimodal study. For mouse tissues, we mapped eQTL results for matching human tissues to the mouse genome, as human and mouse epigenomes show strong conservation^85,86^. We first intersected eQTL variant-gene pairs with peak-gene pairs from the single-cell data and assigned each remaining peak-gene pair the most statistically significant eQTL. We then binned these peak-gene pairs according to the genomic distance between each gene’s TSS and each peak’s center, creating a bin for each of the following ranges of genomic distances: [1 bp, 1 kb), [1 kb, 10kb), [10kb, 100kb), [100kb, 1 Mb). We identified all peak-gene pairs that overlapped significant eQTL variant-gene pairs (*P* < 0.001), and these peak-gene pairs served as the positive examples. To generate a distance-matched set of negative examples, we determined the number of positive examples in each genomic distance bin and selected an equal number of non-significant eQTL variant-gene pairs with the highest *P* values for that bin.

### Chromatin interactions evaluations

We obtained Hi-C data for cell types matching the 10x Multiome, SNARE-seq, and sci-CAR datasets^88–90^. We used 5C data for the SHARE-seq dataset^91^ due to the unavailability of mouse skin Hi-C data. Chromatin interactions were mapped to a peak-gene pair by intersecting the genomic windows in the chromatin interactions data with the peak and the TSS of the gene. For Hi-C data, peak-gene pairs that were associated with the top 1% of chromatin contact frequencies were treated as positive examples, while the remaining peak-gene pairs were considered negative examples. For the 5C dataset, which evaluated the statistical significance of chromatin loops, peak-gene pairs that were associated with significant looping interactions (*P* < 0.001) were treated as positive examples, and the remaining peak-gene pairs were considered negative examples.

### GrID-Net vs. correlation peak-gene prioritization

For the human corticogenesis dataset, we averaged the rankings of all candidate peak-gene pairs across the five trials for each of the GrID-Net, Pearson correlation, and pseudocell correlation methods. We then calculated the difference between GrID-Net’s average rankings for each peak-gene pair with those of the Pearson correlation and pseudocell correlation methods. The peak-gene pairs with the most positive differences represent the peak-gene pairs that GrID-Net prioritizes over the method against which GrID-Net is being compared. To evaluate enrichment of gene sets in peak-gene pairs prioritized by GrID-Net, genes were ranked with respect to these differences and enrichment was computed using GSEA Preranked^34^.

### Enhancer states

We obtained H3K27ac and H3K4me1 ChIP-seq data for all samples of human neural progenitor cells and fetal neural cells in ChIP-Atlas ^30^. We identified statistically significant ChIP-seq peaks (MACS2 *Q* < 10^−10^) that were positioned fully within ATAC-seq peaks from the human corticogenesis dataset. Active enhancers were defined as ATAC-seq peaks containing overlapping H3K4me1 and H3K27ac ChIP-seq peaks, while poised enhancers were defined as ATAC-seq peaks containing H3k4me1 ChIP-seq peaks and no H3K27ac ChIP-seq peaks.

### Genomic distance-dependent GrID-Net scores

We constructed a general model that integrated GrID-Net scores with genomic distance to account for known patterns of genomic distance-dependent transcription factor (TF) regulatory effects^42,43^. The model assumes that TF regulatory effects exponentially decay with increasing genomic distance. To learn the decay rate, we trained a logistic regression model to predict peak-gene pairs overlapping significant skin eQTLs (*P* < 10^−3^) for the SHARE-seq mouse skin dataset.

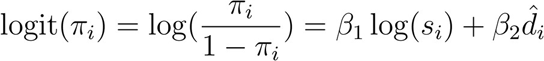

Here, *π*_i_ is the probability of peak-gene pair *i* being a positive example, *s*_*i*_ corresponds to the mean rankings of the GrID-Net scores across trials for the peak-gene pair, and 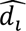 = *max(log_10_ (d_i_) – 40)*, where *d_i_* is the genomic distance in base pairs (bp) between the center of the peak and the TSS of the gene in the peak-gene pair. This formulation assumes that only distal interactions at genomic distances greater than 10 kb are affected by the genome distance-dependent TF regulatory effect.

Positive and negative examples were chosen as described in “eQTL evaluations”. We held out one-third of the example as a test set, and then performed nested 3-fold cross validation on the remaining examples, selecting the model that yielded the highest AUROC across the folds. This model increased the AUROC on the test set compared to the predictions from the averaged GrID-Net rankings (**Figure S3a**). We applied this pre-trained model on the human corticogenesis dataset and found that this model outperformed the baseline predictions from the averaged GrID-Net rankings (**Figure S3b**). These results demonstrated the effectiveness of incorporating genomic distance as an additional feature alongside scores learned from the single-cell multimodal data.

### Linking SCZ GWAS Variants to Genes

We obtained FINEMAP posterior probabilities (*PP*) calculated in the original SCZ GWAS study^41^. We retained all fine-mapped variants with *PP* ≥ 0.05 that overlapped peaks from the human corticogenesis dataset. We identified the peak that matched each SCZ variant and, using the genomic distance-dependent GrID-Net scores, we linked the variant to up to 3 of the peak’s highest scoring genes, retaining only peak-gene pairs in the top 50% of all peak-gene scores.

### Brain-specific gene expression

We obtained consensus tissue-specific transcript levels from the Human Protein Atlas^92,93^. For a given gene, we first calculated the mean and standard deviation of the gene’s transcript levels across all non-brain tissues. The brains-specific tissues were identified to be those from the basal ganglia, cerebellum, cerebral cortex, medulla oblongata, midbrain, pons, and white matter. We then converted the transcript level for the gene in each of the brain-specific tissues into a z-score with respect to the non-brain tissues 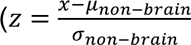, where *x* is the transcript level for the gene in a brain-specific tissue). Genes that had a z-score over 1 in any of the brain-specific tissues were annotated as having brain-specific gene expression.

### TF binding affinity changes

We obtained the reference and SCZ risk alleles for all SCZ variants from the original SCZ GWAS study^41^. We then flanked each of the alleles with the genomic sequences ±20 bp of the SCZ variant position using the GRCh38/hg38 reference genome^94^, generating a reference sequence and SCZ variant sequence. We applied FIMO^95^ to each of these sequences to predict TF binding affinities using position weight matrices (PWMs) from HOCOMOCO v11 (Full Collection)^96^, JASPAR CORE (Non-Redundant) v2^97^, and CIS-BP v2^98^. We evaluated the effect of a specific SCZ variant on a TF’s binding affinity by comparing the FIMO-calculated *p*-values for the TF between the reference sequence and SCZ variant sequence.

### SCZ GPC gene expression

We obtained bulk RNA-seq data from glial progenitor cells (GPCs) produced from 4 SCZ patient-derived induced pluripotent cells (iPSCs) and paired controls subejcts^58,64^, with at least 3 samples per patient. We applied a *log* (1 + *CPM*) transformation to this data.

To evaluate the relationship between TF binding affinity changes caused by SCZ variants to expression changes in GPCs, we first determined peak-gene pairs that are specific to glial differentiation. In the human corticogenesis dataset, we identified genes that were differentially expressed between glial cells and the cycling progenitor cells, which were the root of the corticogenesis trajectory (*P* < 0.05, Welch’s *t*-test). We then used Welch’s *t*-test to compare the expression levels of these glial differentiation-specific genes for each SCZ patient against all of the control samples. For each gene, we identified the most statistically significant difference so as to account for all genes that may be differentially expressed in any of the patients compared to the controls.

### SCZ GPC co-expression pathways

We first identified a set of 26 genes linked to SCZ variants causing at least a 40-fold change in binding affinity for any TF. We identified 3 sets of these SCZ genes that were correlated with each other (Pearson correlation coefficient ≥ 0.6) in the SCZ GPCs and paired controls (SLC12A6, DLX1, DLX2; MFAP4, STAT6, NXPH4; EPN2, P2RX7, ICA1). For each of these sets of SCZ genes, we then determined gene modules that may be implicated in their dysregulation by identifying genes that are highly co-expressed (Pearson correlation coefficient ≥ 0.8) with any of the SCZ genes in the set. We applied gene set enrichment analysis using Enrichr to detect biological processes that were enriched in each of these modules.

---

**Figure S1.**
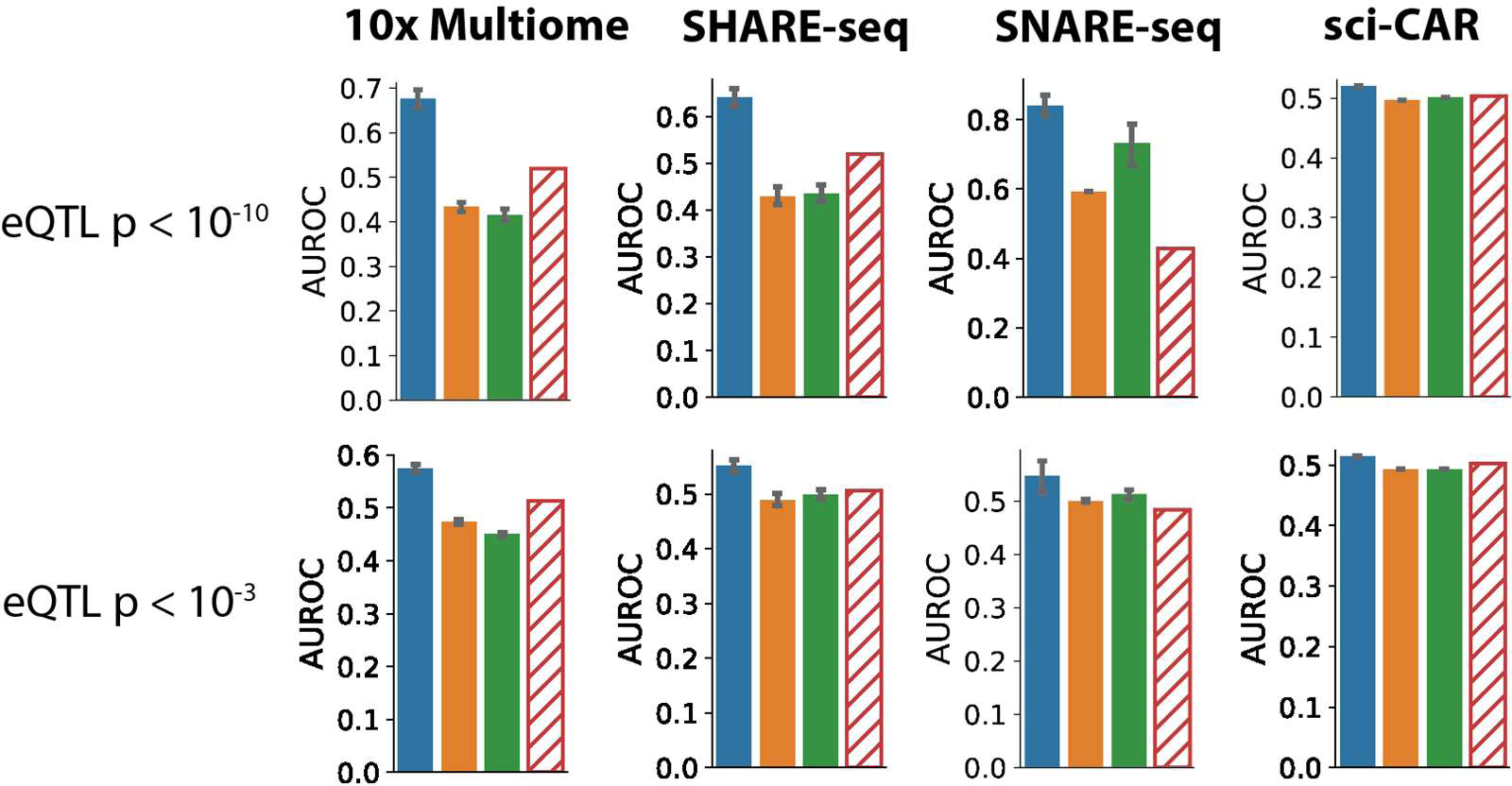
GrID-Net outperforms other methods in eQTL prediction accuracy across different eQTL significance thresholds.

**Figure S2.**
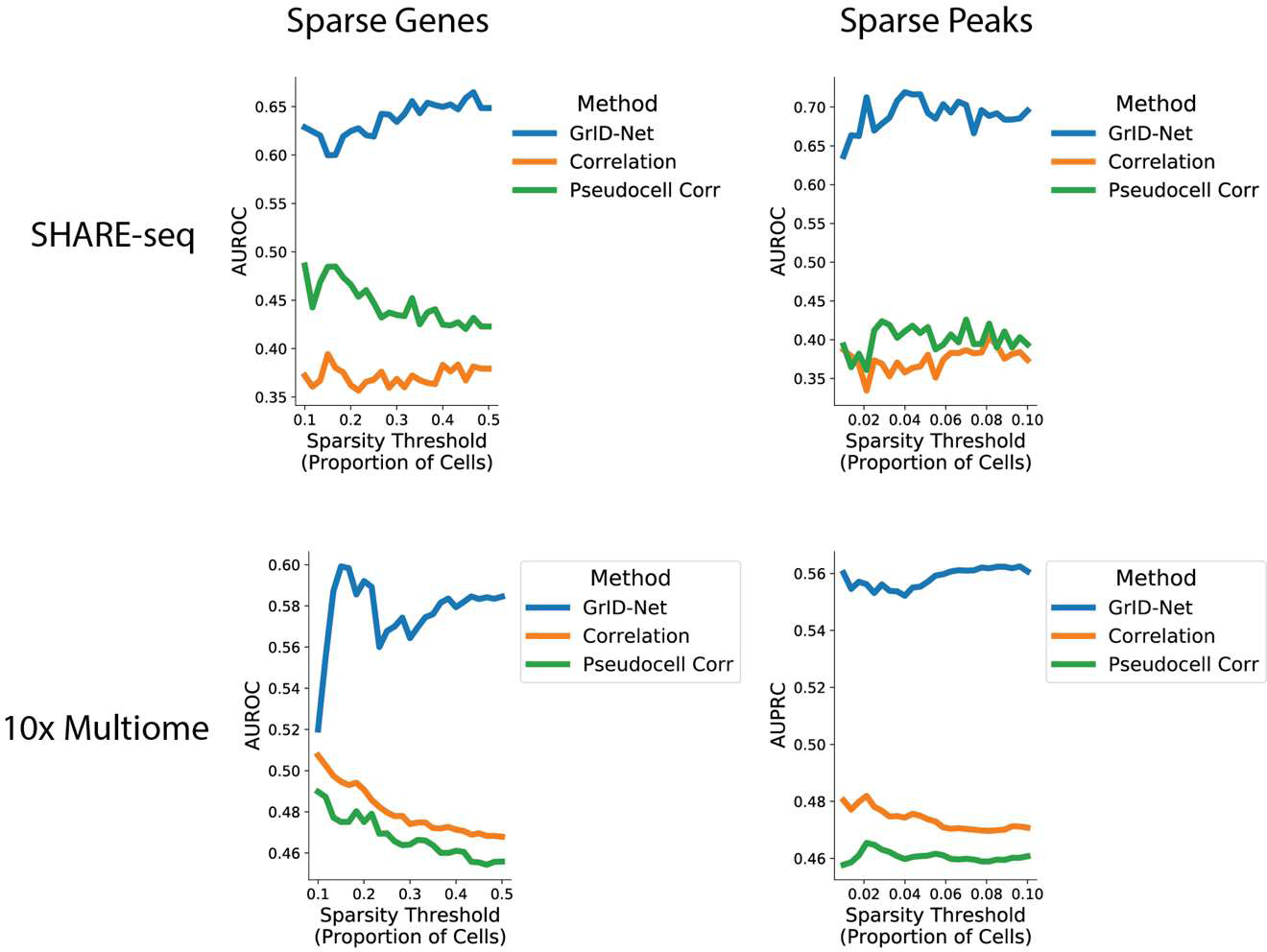
GrID-Net is robust to data sparsity in single-cell datasets.

**Figure S3.**
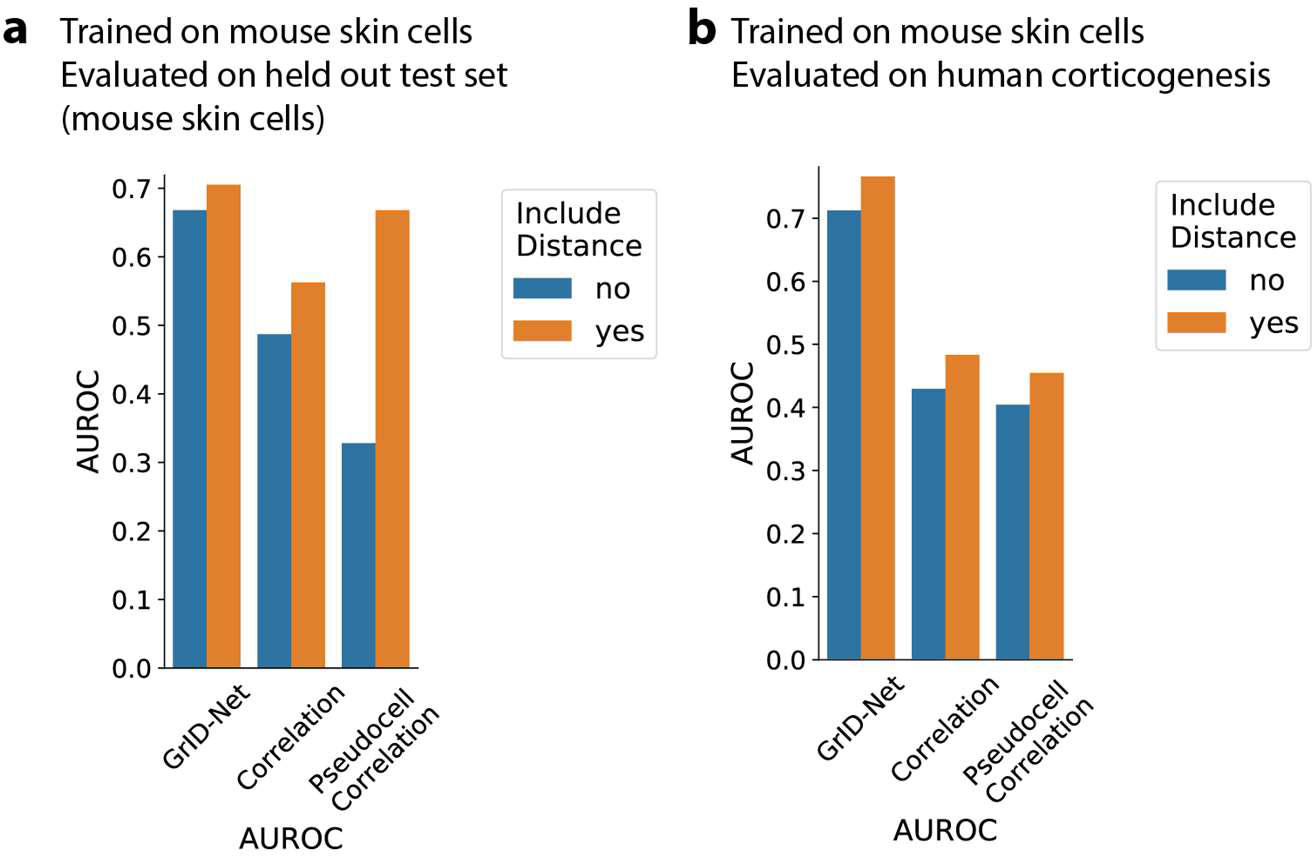
Genomic distance-dependent model improves peak-gene link prediction accuracy (a) within and (b) across species and cell types.

**Figure S4.**
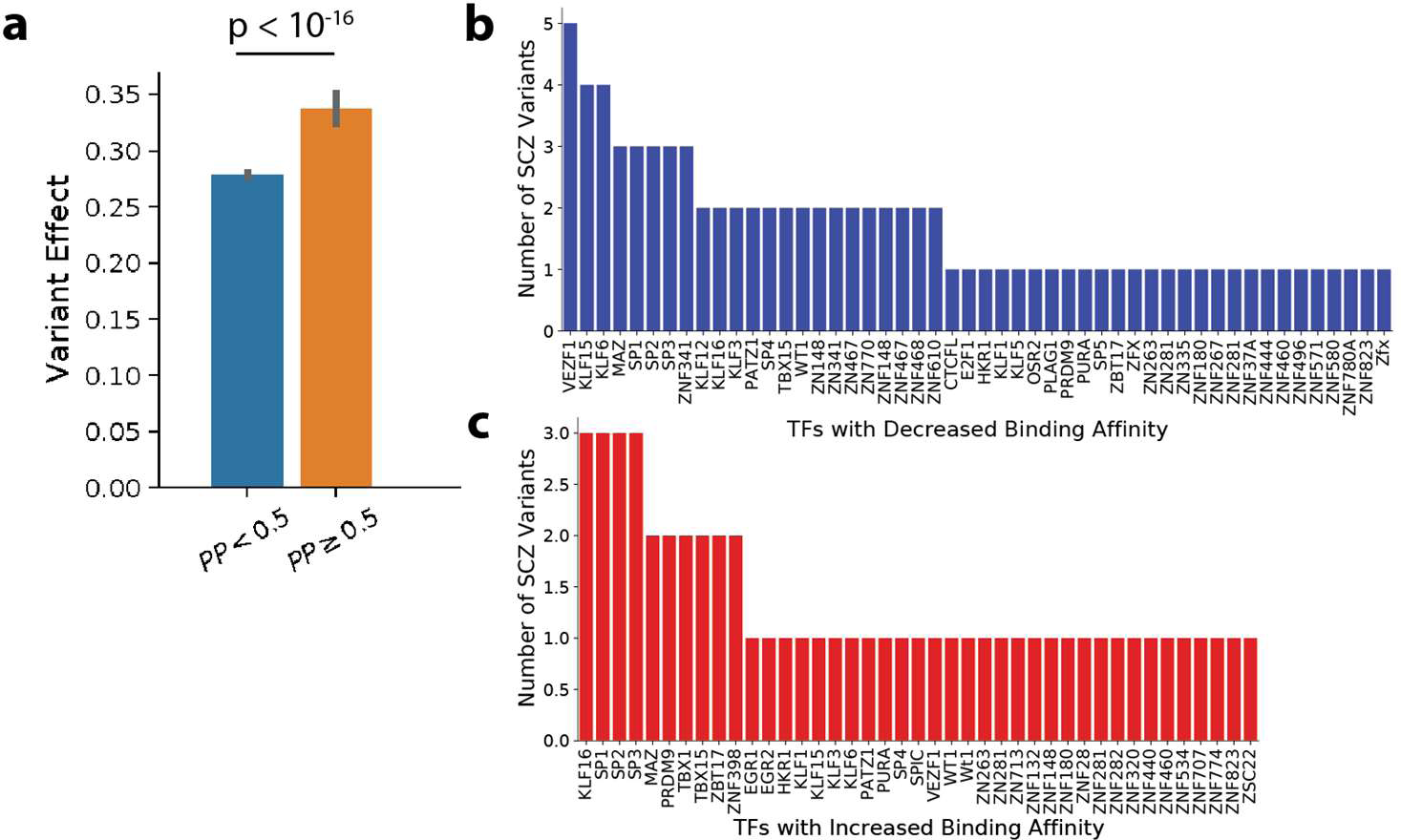
**(a)** Relationship between FINEMAP posterior probabilities (PP) for a SCZ variant and its corresponding TF binding affinity change (Welch’s one-tailed *t*-test). TFs with the most widespread **(b)** increase or **(c)** decrease in binding affinity between the reference and the SCZ risk allele.

## Notes

https://github.com/alexw16/gridnet

### Competing Interest Statement

The authors have declared no competing interest.

### Summary of Updates

Updated exposition

