## Supplementary Note and Tables for "Unveiling causal regulatory mechanisms through cell-state parallax"

### 1 Background on Granger Causality: Standard Formulations

In time series analyses, the classical Granger causality framework uses the vector autoregressive (VAR) model, which can be expressed as follows (Lütkepohl, 2005).

$$y_t = \sum_{\ell=1}^{L} a_1^{(\ell)} y_{t-\ell} + \sum_{\ell=1}^{L} a_2^{(\ell)} x_{t-\ell} + \epsilon_t \tag{1}$$

Here, $y_t, x_t \in \mathbb{R}$ are values at time $t$ for two stationary time series $y$ and $x$, respectively; $a_1^{(\ell)}$ and $a_2^{(\ell)}$ are coefficients that specify how lag $\ell$ affects the future of time series $y$; and $\epsilon_t$ is a zero-mean noise term. In this model, $y_t$ is assumed to be a linear combination of the $L$ most recent values of $y$ and $x$ each, with $x$ said to Granger cause time series $y$ if and only if $a_2^{(\ell)} \neq 0$ for all $\ell$.

A more generalized formulation — the one we follow — for performing Granger causal inference is to consider two related models for forecasting time series $y$. A *full* model considers the past values of both $y$ and $x$ to forecast $y$, while a *reduced* model excludes the effect of $x$, only containing terms related to the past values of $y$.

$$y_t = f^{(full)}(y_{t-1}, ..., y_{t-L}; x_{t-1}, ..., x_{t-L}) + \epsilon_t \tag{2}$$

$$y_t = f^{(reduced)}(y_{t-1}, ..., y_{t-L}) + \epsilon_t \tag{3}$$

Here, $f^{(full)}(\cdot)$ and $f^{(reduced)}(\cdot)$ are generalized functions that specify how the value of $y_t$ depends on past values of $y$ and $x$. The predictions of the two models are then compared, upon which a Granger causal relationship is declared if the full model's predictions are significantly more accurate.

### 2 Granger Causality on a DAG: Graph Neural Network Formulation

Let the data be represented by a DAG $\mathcal{G} = (\mathcal{V}, \mathcal{E})$ with $n = |\mathcal{V}|$ nodes (i.e., observations) and directed edges $\mathcal{E}$ indicating the information flow or partial order between these observations. For instance, when applied to the case of standard Granger causal inference on time series data, $\mathcal{G}$ would be the linear graph corresponding to the time series. Let $\boldsymbol{y}, \boldsymbol{x} \in \mathbb{R}^n$ correspond to the values of variables $y$ and $x$ on the nodes in $\mathcal{V}$, with $x$ putatively Granger-causing $y$. To infer Granger causality, we compare the full and reduced models as above:

$$\boldsymbol{y} = g(\tilde{\boldsymbol{h}}_y^{(full)} + c\tilde{\boldsymbol{h}}_x^{(full)}) + \boldsymbol{\epsilon} \tag{4}$$

$$\boldsymbol{y} = g(\tilde{\boldsymbol{h}}_y^{(reduced)}) + \boldsymbol{\epsilon} \tag{5}$$

Here, $\tilde{\boldsymbol{h}}_y^{(full)}, \tilde{\boldsymbol{h}}_y^{(reduced)}, \tilde{\boldsymbol{h}}_x^{(full)} \in \mathbb{R}^n$ represent the historical information of $y$ or $x$ (as denoted by the subscript) that is aggregated by graph neural network (GNN) layers. For a history $\tilde{\boldsymbol{h}}$, the value $\tilde{\boldsymbol{h}}[v]$ at node $v \in \mathcal{V}$ is the information accumulated from $v$'s ancestors in $\mathcal{G}$. While $\tilde{\boldsymbol{h}}_y^{(full)}$ and $\tilde{\boldsymbol{h}}_y^{(reduced)}$ are outputs determined from similar architectures, their learned weights and bias terms end up being different since $\tilde{\boldsymbol{h}}_y^{(full)}$ is influenced by its interaction with $\tilde{\boldsymbol{h}}_x$. The coefficient $c$ mediates this interaction and describes the effect of $x$'s history on $y$. In addition, $\boldsymbol{\epsilon} \in \mathbb{R}^n$ is a zero-mean noise term. We also include $g(\cdot)$ as an optional link function that maps its input to the support of the distribution of variable $y$. In our single-cell application, we set $g$ as the exponential function since the target variable ($\boldsymbol{y}$) represents normalized transcript counts, which are non-negative.

For brevity, we only describe how $\tilde{\boldsymbol{h}}_x^{(full)}$ is computed, with $\tilde{\boldsymbol{h}}_y^{(full)}$ and $\tilde{\boldsymbol{h}}_y^{(reduced)}$ being independently computed analogously. Also, since $\boldsymbol{x}$ occurs only in the full model, for notational convenience we drop its superscript *(full)* below, writing just $\tilde{\boldsymbol{h}}_x$. We express $\tilde{\boldsymbol{h}}_x$ as the mean of the outputs of $L$ consecutive GNN layers, denoting the layerwise outputs as $\boldsymbol{h}_x^{(\ell)}$:

$$\tilde{\boldsymbol{h}}_x = \frac{1}{L}\sum_{\ell=1}^{L} \boldsymbol{h}_x^{(\ell)} \tag{6}$$

$$\boldsymbol{h}_x^{(\ell)} = \begin{cases} \sigma(w_x^{(\ell)} \boldsymbol{A}_+^T \boldsymbol{h}_x^{(\ell-1)} + b_x^{(\ell)}) & \text{if } \ell > 1 \\ \sigma(w_x^{(\ell)} \boldsymbol{A}^T \boldsymbol{x} + b_x^{(\ell)}) & \text{if } \ell = 1 \end{cases} \tag{7}$$

Here, $w_x^{(\ell)}, b_x^{(\ell)} \in \mathbb{R}$ are the per-layer weight and bias terms, respectively. $\sigma(\cdot)$ represents the nonlinear activation in each of the GNN layers, chosen here to be the hyperbolic tangent function. $\boldsymbol{A}$ and $\boldsymbol{A}_+ \in \mathbb{R}^{n\times n}$ are matrices defined by the DAG structure. $\boldsymbol{A}$ is the column-normalized adjacency matrix of $\mathcal{G}$, with $A_{ij} = \frac{1}{d_j}$ if edge $(i,j) \in \mathcal{E}$ and 0 otherwise, where $d_j$ is the in-degree of node $j$. We note that a DAG does not have self-loops, so the diagonal terms of $\boldsymbol{A}$ are zero. In $\boldsymbol{A}_+$, we include these diagonal terms, setting them to the same value as others in the column (i.e., $d_j$ is incremented by 1 in $\boldsymbol{A}_+$ to preserve normalization).

Together, $\boldsymbol{A}$ and $\boldsymbol{A}_+$ allow us to extend the key intuition of Granger causality to DAGs: predicting a value at node $v$ from the values of its ancestors that are within $L$ steps. The first layer introduces the lag, using information at the parents of $v$ but not the information at $v$ itself. Each subsequent layer introduces the preceding set of parents, with the diagonal term in $\boldsymbol{A}_+$ ensuring that information already stored at $v$ is integrated with the new information propagated via $v$'s parents. The sequence of GNN layers therefore reflects the successive aggregation of past information by traversing $L$ steps backward along the DAG for each node in the graph. Thus, we are able to conceptually match time series-based formulations (e.g., Eqn. 1) but with the crucial advantage of leveraging the richness of a DAG's structure in directing the information flow and aggregating sparse data.

### 3 Comparing Full and Reduced Models to Infer Granger Causality

Let $\Theta_y^{(full)}, \Theta_y^{(reduced)}, \Theta_x^{(full)} \in \mathbb{R}^{2L}$ denote the set of parameters for the full and reduced models, with $\Theta_x^{(full)} = \{w_x^{(1)}, ..., w_x^{(L)}; b_x^{(1)}, ..., b_x^{(L)}\}$ and $\Theta_y^{(full)}, \Theta_y^{(reduced)}$ defined analogously. All the $\Theta$ parameters are jointly learned by minimizing the combined loss of the full and reduced models (Montalto et al., 2015):

$$\mathcal{L}^{(total)} = \sum_{x,y\in\mathcal{P}} \left( \sum_{v\in\mathcal{V}} \mathcal{L}(\hat{y}_v^{(full)}, y_v) + \sum_{v\in\mathcal{V}} \mathcal{L}(\hat{y}_v^{(reduced)}, y_v) \right) \tag{8}$$

where $\hat{y}_v^{(full)}$ and $\hat{y}_v^{(reduced)}$ correspond to the predictions for observation $v$ by the full and reduced models, respectively. We note that the full and reduced models have completely separate parameters. Also, in a multivariate system, not all pairwise combinations of variables may be relevant: $\mathcal{P}$ is the subset of $x, y$ variable pairs whose putative Granger-causal interactions are of interest.

To infer Granger causal interactions between $y$ and $x$, we compare the set of loss terms associated with the full model $\mathcal{L}(\hat{y}_v^{(full)}, y_v)$ to those of the reduced model $\mathcal{L}(\hat{y}_v^{(reduced)}, y_v)$, noting that the precise functional form of $\mathcal{L}$ depends on the domain. If $x$ does not Granger cause $y$, the full and reduced models will have similar explanatory power and hence similar loss values. Otherwise, the full model will have a lower loss. We assess this by a one-tailed F-test, comparing the residual sum of squares of the reduced and full models. We rank the set of candidate $x$–$y$ interactions by the F-statistic, with a higher score corresponding to stronger evidence for a Granger causal interaction.

### 4 Domain-specific Model Customization

The loss $\mathcal{L}$ should be chosen as appropriate for the domain. In our single-cell context, $y$ corresponds to gene expression. We preprocessed RNA-seq transcript counts, normalizing and log transforming them so that the mean squared error loss was appropriate; we note that most methods based on Pearson correlation of gene expression also seek to minimize squared loss, explicitly or implicitly.

The GrID-Net model can also be customized to account for different lags and lookbacks. The number of GNN layers $L$ corresponds to the maximum amount of past information desired. Similarly, to introduce a $k$-hop lag, the first $k$ GNN layers would use the matrix $\boldsymbol{A}$ while the later layers would use $\boldsymbol{A}_{+}$. In the sections above, we described a one-hop lag that we have used for all analyses here.

| | A | B | C | D | E | F | G | H | I |
|---|---|---|---|---|---|---|---|---|---|
| 1 | Table S1 | | | | | | | | |
| 2 | Notes: 10x Multiome data from Trevino et al (PMID: 34390642) | | | | | | | | |
| 3 | Notes: GWAS data from Trubetskoy et al. (PMID: 35396580). Max 3 genes per peak, with 0 being strongest. | | | | | | | | |
| 4 | | | | | | | | | |
| 5 | atac_id_PMID_34390642 | rsID_PMID_35396580 | position | gwas_pp_PMID_35396580 | gridnet_associated_gene | variant_dist_from_TSS | gridnet_score | peak-specfic_rank | nearest_gene_to_variant |
| 6 | 739 | rs6673661 | chr1-2441515 | 0.186 | PLCH2 | 34729 | 0.542570487 | 0 | PEX10 |
| 7 | 739 | rs6673661 | chr1-2441515 | 0.186 | PEX10 | 28546 | 0.538469099 | 1 | PEX10 |
| 8 | 739 | rs6673661 | chr1-2441515 | 0.186 | MORN1 | 51064 | 0.533158197 | 2 | PEX10 |
| 9 | 740 | rs6673880 | chr1-2441729 | 0.202 | PLCH2 | 34119 | 0.541885623 | 0 | PEX10 |
| 10 | 740 | rs6673880 | chr1-2441729 | 0.202 | PEX10 | 29156 | 0.541201119 | 1 | PEX10 |
| 11 | 740 | rs6673880 | chr1-2441729 | 0.202 | MORN1 | 51674 | 0.533515977 | 2 | PEX10 |
| 12 | 742 | rs4648844 | chr1-2443319 | 0.105 | PLCH2 | 32577 | 0.547559269 | 0 | PEX10 |
| 13 | 742 | rs4648844 | chr1-2443319 | 0.105 | PEX10 | 30698 | 0.546243547 | 1 | PEX10 |
| 14 | 742 | rs4648844 | chr1-2443319 | 0.105 | MORN1 | 53216 | 0.538934316 | 2 | PEX10 |
| 15 | 744 | rs6687012 | chr1-2444405 | 0.07 | PLCH2 | 31423 | 0.546202516 | 0 | PEX10 |
| 16 | 744 | rs6687012 | chr1-2444405 | 0.07 | PEX10 | 31852 | 0.541857381 | 1 | PEX10 |
| 17 | 744 | rs6687012 | chr1-2444405 | 0.07 | MORN1 | 54370 | 0.532667588 | 2 | PEX10 |
| 18 | 8248 | rs12092160 | chr1-28824146 | 0.077 | TAF12 | 181087 | 0.516259 | 0 | OPRD1 |
| 19 | 12113 | rs11210892 | chr1-43634413 | 0.334 | KDM4A-AS1 | 72876 | 0.531369976 | 0 | KDM4A |
| 20 | 12113 | rs11210892 | chr1-43634413 | 0.334 | ST3GAL3-AS1 | 92850 | 0.527466182 | 1 | KDM4A |
| 21 | 12113 | rs11210892 | chr1-43634413 | 0.334 | HYI | 180185 | 0.52307496 | 2 | KDM4A |
| 22 | 32210 | rs72749142 | chr1-200982179 | 0.215 | KIF21B | 23131 | 0.514867079 | 0 | KIF21B |
| 23 | 32220 | rs12126806 | chr1-200994697 | 0.339 | KIF21B | 10432 | 0.530049201 | 0 | KIF21B |
| 24 | 32232 | rs12122721 | chr1-201015352 | 0.114 | KIF21B | 9698 | 0.524731132 | 0 | KIF21B |

| | A | B | C | D | E | F | G | H | I |
|---|---|---|---|---|---|---|---|---|---|
| 25 | 40469 | rs72759887 | chr1-243457054 | 0.174 | SDCCAG8 | 30562 | 0.515256445 | 0 | SDCCAG8 |
| 26 | 41977 | rs17247190 | chr2-2312029 | 0.058 | MYT1L-AS1 | 6893 | 0.578665419 | 0 | MYT1L-AS1 |
| 27 | 41981 | rs10206268 | chr2-2322151 | 0.1 | MYT1L-AS1 | 2432 | 0.572811384 | 0 | MYT1L-AS1 |
| 28 | 41985 | rs13419191 | chr2-2325568 | 0.078 | MYT1L-AS1 | 6159 | 0.579181682 | 0 | MYT1L |
| 29 | 41986 | rs6708950 | chr2-2326346 | 0.084 | MYT1L-AS1 | 6724 | 0.579964542 | 0 | MYT1L |
| 30 | 41991 | rs3749051 | chr2-2331105 | 0.074 | MYT1L-AS1 | 11414 | 0.575587953 | 0 | MYT1L |
| 31 | 46895 | rs527882868 | chr2-25216094 | 0.134 | ARNILA | 159381 | 0.532235201 | 0 | LINC01381 |
| 32 | 46895 | rs527882868 | chr2-25216094 | 0.134 | DTNB-AS1 | 204653 | 0.529724605 | 1 | LINC01381 |
| 33 | 46901 | rs145260949 | chr2-25230278 | 0.375 | ARNILA | 145211 | 0.529496445 | 0 | LINC01381 |
| 34 | 46901 | rs145260949 | chr2-25230278 | 0.375 | DTNB-AS1 | 190483 | 0.526417128 | 1 | LINC01381 |
| 35 | 46901 | rs145260949 | chr2-25230278 | 0.375 | DNAJC27-AS1 | 228718 | 0.51076429 | 2 | LINC01381 |
| 36 | 46904 | rs114568253 | chr2-25237494 | 0.698 | ARNILA | 138241 | 0.532404939 | 0 | LINC01381 |
| 37 | 46904 | rs114568253 | chr2-25237494 | 0.698 | DTNB-AS1 | 183513 | 0.527613766 | 1 | LINC01381 |
| 38 | 46904 | rs114568253 | chr2-25237494 | 0.698 | DNAJC27-AS1 | 235688 | 0.515999658 | 2 | LINC01381 |
| 39 | 48820 | rs3770752 | chr2-37348993 | 0.289 | QPCT | 18989 | 0.575132938 | 0 | QPCT |
| 40 | 48820 | rs3770752 | chr2-37348993 | 0.289 | CEBPZOS | 135325 | 0.53993796 | 1 | QPCT |
| 41 | 48820 | rs3770752 | chr2-37348993 | 0.289 | CEBPZ | 117423 | 0.515244683 | 2 | QPCT |
| 42 | 54931 | rs6722190 | chr2-72941464 | 0.217 | EMX1 | 8125 | 0.527663108 | 0 | EMX1 |
| 43 | 65128 | rs12991836 | chr2-144383974 | 0.723 | ZEB2-AS1 | 136397 | 0.526894864 | 0 | GTDC1 |

| | A | B | C | D | E | F | G | H | I |
|---|---|---|---|---|---|---|---|---|---|
| 44 | 65128 | rs12991836 | chr2-144383974 | 0.723 | GTDC1 | 51187 | 0.508338935 | 1 | GTDC1 |
| 45 | 67955 | rs11693702 | chr2-161945674 | 0.061 | DPP4 | 128238 | 0.531599291 | 0 | DPP4 |
| 46 | 67959 | rs7604885 | chr2-161949898 | 0.065 | DPP4 | 123809 | 0.533598168 | 0 | DPP4 |
| 47 | 69750 | rs62183854 | chr2-172090705 | 0.098 | DLX2 | 11880 | 0.574525291 | 0 | DLX1 |
| 48 | 69750 | rs62183854 | chr2-172090705 | 0.098 | DLX1 | 3153 | 0.573090269 | 1 | DLX1 |
| 49 | 69750 | rs62183854 | chr2-172090705 | 0.098 | METAP1D | 19857 | 0.510134481 | 2 | DLX1 |
| 50 | 69751 | rs62183855 | chr2-172091721 | 0.092 | DLX2 | 10725 | 0.561547164 | 0 | DLX1 |
| 51 | 69751 | rs62183855 | chr2-172091721 | 0.092 | DLX1 | 4308 | 0.560724579 | 1 | DLX1 |
| 52 | 69752 | rs12469319 | chr2-172092378 | 0.075 | DLX2 | 10096 | 0.577122362 | 0 | DLX1 |
| 53 | 69752 | rs12469319 | chr2-172092378 | 0.075 | DLX1 | 4937 | 0.570485121 | 1 | DLX1 |
| 54 | 69752 | rs12469319 | chr2-172092378 | 0.075 | METAP1D | 21641 | 0.507753565 | 2 | DLX1 |
| 55 | 69768 | rs62183859 | chr2-172104047 | 0.059 | DLX2 | 886 | 0.5683438 | 0 | DLX2-DT |
| 56 | 69768 | rs62183859 | chr2-172104047 | 0.059 | DLX1 | 16419 | 0.556855484 | 1 | DLX2-DT |
| 57 | 69773 | rs62184960 | chr2-172110061 | 0.251 | DLX2 | 6861 | 0.583559414 | 0 | DLX2-DT |
| 58 | 69773 | rs62184960 | chr2-172110061 | 0.251 | DLX1 | 22394 | 0.560874212 | 1 | DLX2-DT |
| 59 | 74180 | rs6736049 | chr2-200350990 | 1 | KCTD18 | 158842 | 0.509115424 | 0 | SPATS2L |
| 60 | 87494 | rs4293666 | chr3-30029954 | 0.063 | RBMS3 | 41656 | 0.525636035 | 0 | RBMS3-AS1 |

| | A | B | C | D | E | F | G | H | I |
|---|---|---|---|---|---|---|---|---|---|
| 61 | 90948 | rs2014830 | chr3-50134964 | 0.993 | SEMA3F-AS1 | 20645 | 0.534867835 | 0 | SEMA3F-AS1 |
| 62 | 90948 | rs2014830 | chr3-50134964 | 0.993 | RBM5-AS1 | 33887 | 0.527057431 | 1 | SEMA3F-AS1 |
| 63 | 90948 | rs2014830 | chr3-50134964 | 0.993 | SLC38A3 | 82542 | 0.521447309 | 2 | SEMA3F-AS1 |
| 64 | 91067 | rs709210 | chr3-50320438 | 0.609 | TUSC2 | 7530 | 0.562275017 | 0 | HYAL2 |
| 65 | 91067 | rs709210 | chr3-50320438 | 0.609 | HYAL2 | 484 | 0.55831949 | 1 | HYAL2 |
| 66 | 91067 | rs709210 | chr3-50320438 | 0.609 | CYB561D2 | 30532 | 0.552351806 | 2 | HYAL2 |
| 67 | 91647 | rs1108842 | chr3-52686064 | 0.076 | GLT8D1 | 19826 | 0.567387528 | 0 | GNL3 |
| 68 | 91647 | rs1108842 | chr3-52686064 | 0.076 | NEK4 | 84733 | 0.541165287 | 1 | GNL3 |
| 69 | 91647 | rs1108842 | chr3-52686064 | 0.076 | SMIM4 | 149101 | 0.518793056 | 2 | GNL3 |
| 70 | 93934 | rs59971314 | chr3-63928942 | 0.058 | THOC7 | 88731 | 0.53377672 | 0 | SCAANT1 |
| 71 | 93934 | rs59971314 | chr3-63928942 | 0.058 | PSMD6-AS2 | 75012 | 0.52715185 | 1 | SCAANT1 |
| 72 | 93934 | rs59971314 | chr3-63928942 | 0.058 | C3orf49 | 105010 | 0.527041328 | 2 | SCAANT1 |
| 73 | 110066 | rs9825834 | chr3-177049320 | 0.061 | TBL1XR1-AS1 | 3998 | 0.570394935 | 0 | TBL1XR1-AS1 |
| 74 | 146505 | rs4132385 | chr5-61319052 | 0.054 | SMIM15-AS1 | 156789 | 0.515146482 | 0 | ZSWIM6 |
| 75 | 146505 | rs4132385 | chr5-61319052 | 0.054 | SMIM15 | 156391 | 0.508482321 | 1 | ZSWIM6 |
| 76 | 146522 | rs4604142 | chr5-61346768 | 0.097 | SMIM15-AS1 | 184343 | 0.51951195 | 0 | ZSWIM6 |
| 77 | 146522 | rs4604142 | chr5-61346768 | 0.097 | SMIM15 | 183945 | 0.516539263 | 1 | ZSWIM6 |

| | A | B | C | D | E | F | G | H | I |
|---|---|---|---|---|---|---|---|---|---|
| 78 | 158595 | rs10447226 | chr5-139649225 | 0.056 | PSD2 | 172484 | 0.511470857 | 0 | CXXC5-AS1 |
| 79 | 158635 | rs2336880 | chr5-139684143 | 0.109 | PSD2 | 137581 | 0.524334139 | 0 | CXXC5-AS1 |
| 80 | 158636 | rs10059371 | chr5-139684682 | 0.069 | PSD2 | 137020 | 0.519555187 | 0 | CXXC5-AS1 |
| 81 | 158637 | rs34510678 | chr5-139685250 | 0.11 | PSD2 | 136407 | 0.518737374 | 0 | CXXC5-AS1 |
| 82 | 158638 | rs35123781 | chr5-139685595 | 0.074 | PSD2 | 135878 | 0.51544178 | 0 | CXXC5-AS1 |
| 83 | 158639 | rs9687282 | chr5-139686403 | 0.132 | PSD2 | 135231 | 0.523296394 | 0 | CXXC5-AS1 |
| 84 | 181358 | rs76593596 | chr6-96016752 | 0.056 | FUT9 | 498 | 0.535817862 | 0 | FUT9 |
| 85 | 183445 | rs9374040 | chr6-108676232 | 0.053 | FOXO3 | 19440 | 0.513351999 | 0 | LINC00222 |
| 86 | 189068 | rs2206956 | chr6-146418092 | 0.232 | RAB32 | 125455 | 0.525876397 | 0 | RAB32 |
| 87 | 189071 | rs2300627 | chr6-146421903 | 0.112 | RAB32 | 121535 | 0.527661426 | 0 | RAB32 |
| 88 | 196755 | rs11972718 | chr7-8509557 | 0.395 | ICA1 | 246939 | 0.50544895 | 0 | NXPH1 |
| 89 | 196757 | rs6978582 | chr7-8511018 | 0.097 | ICA1 | 248372 | 0.512251137 | 0 | NXPH1 |
| 90 | 224921 | rs3808581 | chr8-26392531 | 0.146 | BNIP3L | 2729 | 0.540524783 | 0 | BNIP3L |
| 91 | 225237 | rs2472554 | chr8-27470837 | 0.134 | EPHX2 | 44477 | 0.560356245 | 0 | CHRNA2 |
| 92 | 236502 | rs4734654 | chr8-102657763 | 0.154 | KLF10 | 3121 | 0.571043948 | 0 | KLF10 |
| 93 | 243774 | rs10092551 | chr8-143793775 | 0.166 | IQANK1 | 4146 | 0.572068173 | 0 | SCRIB |
| 94 | 243774 | rs10092551 | chr8-143793775 | 0.166 | SCRIB | 14516 | 0.566340553 | 1 | SCRIB |
| 95 | 243774 | rs10092551 | chr8-143793775 | 0.166 | ZNF707 | 85610 | 0.5282787 | 2 | SCRIB |

| | A | B | C | D | E | F | G | H | I |
|---|---|---|---|---|---|---|---|---|---|
| 96 | 243775 | rs11993154 | chr8-143794599 | 0.059 | IQANK1 | 5022 | 0.566030711 | 0 | SCRIB |
| 97 | 243775 | rs11993154 | chr8-143794599 | 0.059 | SCRIB | 13640 | 0.555824944 | 1 | SCRIB |
| 98 | 243775 | rs11993154 | chr8-143794599 | 0.059 | ZNF707 | 86486 | 0.517503918 | 2 | SCRIB |
| 99 | 243799 | rs55930529 | chr8-143823571 | 0.155 | IQANK1 | 33825 | 0.563010124 | 0 | PUF60 |
| 100 | 243799 | rs55930529 | chr8-143823571 | 0.155 | SCRIB | 14663 | 0.551028094 | 1 | PUF60 |
| 101 | 243799 | rs55930529 | chr8-143823571 | 0.155 | NRBP2 | 16790 | 0.526679617 | 2 | PUF60 |
| 102 | 249506 | rs9644880 | chr9-36318848 | 0.249 | GNE | 41592 | 0.548573002 | 0 | GNE |
| 103 | 251091 | rs500102 | chr9-74743829 | 0.271 | RORB-AS1 | 244279 | 0.533410233 | 0 | TRPM6 |
| 104 | 251091 | rs500102 | chr9-74743829 | 0.271 | TRPM6 | 143814 | 0.516731714 | 1 | TRPM6 |
| 105 | 251091 | rs500102 | chr9-74743829 | 0.271 | C9orf40 | 209006 | 0.508641888 | 2 | TRPM6 |
| 106 | 254204 | rs3815563 | chr9-93450864 | 0.076 | FAM120A OS | 148 | 0.563113902 | 0 | FAM120A |
| 107 | 254204 | rs3815563 | chr9-93450864 | 0.076 | C9orf129 | 104479 | 0.533813833 | 1 | FAM120A |
| 108 | 254204 | rs3815563 | chr9-93450864 | 0.076 | PHF2 | 125496 | 0.506014514 | 2 | FAM120A |
| 109 | 254219 | rs7048853 | chr9-93490896 | 0.123 | C9orf129 | 144475 | 0.531655177 | 0 | FAM120A OS |
| 110 | 254219 | rs7048853 | chr9-93490896 | 0.123 | FAM120A OS | 39644 | 0.530378251 | 1 | FAM120A OS |
| 111 | 254219 | rs7048853 | chr9-93490896 | 0.123 | PHF2 | 85500 | 0.507471776 | 2 | FAM120A OS |
| 112 | 254223 | rs59582399 | chr9-93523881 | 0.072 | FAM120A OS | 72777 | 0.521144356 | 0 | PHF2 |

| | A | B | C | D | E | F | G | H | I |
|---|---|---|---|---|---|---|---|---|---|
| 113 | 254223 | rs59582399 | chr9-93523881 | 0.072 | C9orf129 | 177608 | 0.519605621 | 1 | PHF2 |
| 114 | 254223 | rs59582399 | chr9-93523881 | 0.072 | PHF2 | 52367 | 0.513118383 | 2 | PHF2 |
| 115 | 262121 | rs77206190 | chr9-132017990 | 0.052 | NTNG2 | 143463 | 0.524969583 | 0 | MED27 |
| 116 | 281589 | rs10883761 | chr10-102660261 | 0.052 | SFXN2 | 65842 | 0.531024866 | 0 | TRIM8 |
| 117 | 281589 | rs10883761 | chr10-102660261 | 0.052 | WBP1L | 172063 | 0.528737152 | 1 | TRIM8 |
| 118 | 281589 | rs10883761 | chr10-102660261 | 0.052 | C10orf95-AS1 | 206318 | 0.516344221 | 2 | TRIM8 |
| 119 | 281592 | rs4244354 | chr10-102666420 | 0.061 | SFXN2 | 60003 | 0.527725981 | 0 | TRIM8 |
| 120 | 281592 | rs4244354 | chr10-102666420 | 0.061 | WBP1L | 166224 | 0.521215382 | 1 | TRIM8 |
| 121 | 281592 | rs4244354 | chr10-102666420 | 0.061 | C10orf95-AS1 | 212157 | 0.512767191 | 2 | TRIM8 |
| 122 | 299026 | rs58950470 | chr11-65616284 | 0.167 | MAP3K11 | 2012 | 0.562014346 | 0 | PCNX3 |
| 123 | 299026 | rs58950470 | chr11-65616284 | 0.167 | PCNX3 | 13663 | 0.555640335 | 1 | PCNX3 |
| 124 | 299026 | rs58950470 | chr11-65616284 | 0.167 | RNASEH2C | 104050 | 0.544327436 | 2 | PCNX3 |
| 125 | 307188 | rs7112715 | chr11-113532053 | 0.499 | TMPRSS5 | 174149 | 0.522753169 | 0 | DRD2 |
| 126 | 307188 | rs7112715 | chr11-113532053 | 0.499 | TTC12 | 215485 | 0.510079731 | 1 | DRD2 |
| 127 | 307201 | rs7928017 | chr11-113578040 | 0.146 | TMPRSS5 | 127878 | 0.518010718 | 0 | DRD2 |
| 128 | 307201 | rs7928017 | chr11-113578040 | 0.146 | ZW10 | 195240 | 0.513555678 | 1 | DRD2 |
| 129 | 310377 | rs7927437 | chr11-123525279 | 0.134 | ZNF202 | 216371 | 0.514258875 | 0 | GRAMD1B |

| | A | B | C | D | E | F | G | H | I |
|---|---|---|---|---|---|---|---|---|---|
| 130 | 312920 | rs502834 | chr11-133947438 | 0.08 | IGSF9B | 8990 | 0.543228502 | 0 | IGSF9B |
| 131 | 312949 | rs4936216 | chr11-133983113 | 0.097 | IGSF9B | 26101 | 0.529105423 | 0 | IGSF9B |
| 132 | 312971 | rs78971498 | chr11-134014628 | 0.108 | IGSF9B | 57698 | 0.514349952 | 0 | LINC02730 |
| 133 | 312971 | rs78971498 | chr11-134014628 | 0.108 | ACAD8 | 247233 | 0.505918393 | 1 | LINC02730 |
| 134 | 312971 | rs78971498 | chr11-134014628 | 0.108 | THYN1 | 238349 | 0.505590542 | 2 | LINC02730 |
| 135 | 312974 | rs58406308 | chr11-134023085 | 0.07 | THYN1 | 230134 | 0.519307004 | 0 | LINC02730 |
| 136 | 312974 | rs58406308 | chr11-134023085 | 0.07 | ACAD8 | 239018 | 0.516656094 | 1 | LINC02730 |
| 137 | 312974 | rs58406308 | chr11-134023085 | 0.07 | IGSF9B | 65913 | 0.515617228 | 2 | LINC02730 |
| 138 | 312980 | rs7938834 | chr11-134029834 | 0.055 | IGSF9B | 72631 | 0.508912703 | 0 | LINC02730 |
| 139 | 312980 | rs7938834 | chr11-134029834 | 0.055 | THYN1 | 223416 | 0.505300548 | 1 | LINC02730 |
| 140 | 312985 | rs2236657 | chr11-134034290 | 0.073 | THYN1 | 218901 | 0.516231899 | 0 | LINC02731 |
| 141 | 312985 | rs2236657 | chr11-134034290 | 0.073 | IGSF9B | 77146 | 0.512657549 | 1 | LINC02731 |
| 142 | 312985 | rs2236657 | chr11-134034290 | 0.073 | ACAD8 | 227785 | 0.511137995 | 2 | LINC02731 |
| 143 | 312987 | rs687187 | chr11-134035439 | 0.083 | IGSF9B | 78635 | 0.51213344 | 0 | LINC02731 |
| 144 | 312987 | rs687187 | chr11-134035439 | 0.083 | THYN1 | 217412 | 0.509856118 | 1 | LINC02731 |
| 145 | 313082 | rs11223774 | chr11-134377421 | 0.394 | GLB1L2 | 45508 | 0.540910206 | 0 | B3GAT1 |
| 146 | 313082 | rs11223774 | chr11-134377421 | 0.394 | THYN1 | 124081 | 0.521707493 | 1 | B3GAT1 |

| | A | B | C | D | E | F | G | H | I |
|---|---|---|---|---|---|---|---|---|---|
| 147 | 313082 | rs11223774 | chr11-134377421 | 0.394 | ACAD8 | 115197 | 0.52013415 | 2 | B3GAT1 |
| 148 | 313823 | rs11062170 | chr12-2239678 | 0.095 | DCP1B | 234981 | 0.516765514 | 0 | CACNA1C-AS4 |
| 149 | 320649 | rs706790 | chr12-50070687 | 0.07 | ASIC1 | 9584 | 0.539298286 | 0 | ASIC1 |
| 150 | 320649 | rs706790 | chr12-50070687 | 0.07 | COX14 | 41323 | 0.523805669 | 1 | ASIC1 |
| 151 | 320649 | rs706790 | chr12-50070687 | 0.07 | NCKAP5L | 241664 | 0.523190087 | 2 | ASIC1 |
| 152 | 322233 | rs324017 | chr12-57094031 | 0.962 | NAB2 | 1695 | 0.561457227 | 0 | NAB2 |
| 153 | 322233 | rs324017 | chr12-57094031 | 0.962 | STAT6 | 17069 | 0.557972495 | 1 | NAB2 |
| 154 | 322233 | rs324017 | chr12-57094031 | 0.962 | NXPH4 | 125525 | 0.51878358 | 2 | NAB2 |
| 155 | 329143 | rs4246263 | chr12-104243799 | 0.054 | EID3 | 59709 | 0.54544124 | 0 | TXNRD1 |
| 156 | 329143 | rs4246263 | chr12-104243799 | 0.054 | NFYB | 105289 | 0.520740127 | 1 | TXNRD1 |
| 157 | 329143 | rs4246263 | chr12-104243799 | 0.054 | GLT8D2 | 193394 | 0.514301973 | 2 | TXNRD1 |
| 158 | 332744 | rs77278225 | chr12-120844458 | 0.287 | C12orf43 | 171790 | 0.511160103 | 0 | SPPL3 |
| 159 | 332751 | rs7486605 | chr12-120903362 | 0.981 | C12orf43 | 112918 | 0.51529939 | 0 | SPPL3 |
| 160 | 332751 | rs7486605 | chr12-120903362 | 0.981 | P2RX7 | 229409 | 0.509123176 | 1 | SPPL3 |
| 161 | 333161 | rs1979237 | chr12-122841348 | 1 | CCDC62 | 52548 | 0.530261305 | 0 | HIP1R |
| 162 | 340007 | rs953102 | chr13-43699249 | 0.1 | CCDC122 | 180095 | 0.521411178 | 0 | ENOX1 |
| 163 | 364705 | rs11847162 | chr14-103346720 | 0.112 | EIF5 | 5489 | 0.50563941 | 0 | EIF5 |

| | A | B | C | D | E | F | G | H | I |
|---|---|---|---|---|---|---|---|---|---|
| 164 | 367234 | rs117799466 | chr15-34367316 | 0.906 | SLC12A6 | 29263 | 0.545975566 | 0 | LPCAT4 |
| 165 | 367234 | rs117799466 | chr15-34367316 | 0.906 | LPCAT4 | 24 | 0.537373474 | 1 | LPCAT4 |
| 166 | 367234 | rs117799466 | chr15-34367316 | 0.906 | NOP10 | 23869 | 0.53358522 | 2 | LPCAT4 |
| 167 | 368375 | rs56205728 | chr15-40275036 | 0.543 | PAK6 | 692 | 0.579155222 | 0 | ANKRD63 |
| 168 | 368375 | rs56205728 | chr15-40275036 | 0.543 | INAFM2 | 50094 | 0.537601762 | 1 | ANKRD63 |
| 169 | 368375 | rs56205728 | chr15-40275036 | 0.543 | BMF | 165744 | 0.524451257 | 2 | ANKRD63 |
| 170 | 377575 | rs7178152 | chr15-89243824 | 0.152 | RLBP1 | 22070 | 0.541695045 | 0 | FANCI |
| 171 | 377660 | rs208827 | chr15-89399100 | 0.066 | LINC0092 8 | 124676 | 0.529049087 | 0 | MIR9-3HG |
| 172 | 377660 | rs208827 | chr15-89399100 | 0.066 | RLBP1 | 177268 | 0.527834436 | 1 | MIR9-3HG |
| 173 | 377660 | rs208827 | chr15-89399100 | 0.066 | POLG | 64429 | 0.526208416 | 2 | MIR9-3HG |
| 174 | 377662 | rs182317 | chr15-89400370 | 0.055 | POLG | 65737 | 0.530252145 | 0 | MIR9-3HG |
| 175 | 377662 | rs182317 | chr15-89400370 | 0.055 | RLBP1 | 178576 | 0.521037796 | 1 | MIR9-3HG |
| 176 | 377662 | rs182317 | chr15-89400370 | 0.055 | LINC0092 8 | 123368 | 0.513989729 | 2 | MIR9-3HG |
| 177 | 378059 | rs4702 | chr15-90883330 | 0.999 | MAN2A2 | 20408 | 0.571502915 | 0 | FES |
| 178 | 378059 | rs4702 | chr15-90883330 | 0.999 | FURIN | 1867 | 0.558483091 | 1 | FES |
| 179 | 378059 | rs4702 | chr15-90883330 | 0.999 | UNC45A | 46398 | 0.553085894 | 2 | FES |
| 180 | 396302 | rs4790338 | chr17-1371519 | 0.095 | INPP5K | 144916 | 0.518666926 | 0 | YWHAE |
| 181 | 396302 | rs4790338 | chr17-1371519 | 0.095 | MYO1C | 119492 | 0.5175162 | 1 | YWHAE |

| | A | B | C | D | E | F | G | H | I |
|---|---|---|---|---|---|---|---|---|---|
| 182 | 396302 | rs4790338 | chr17-1371519 | 0.095 | PITPNA-AS1 | 145186 | 0.507254385 | 2 | YWHAE |
| 183 | 396316 | rs28365859 | chr17-1400484 | 0.165 | INPP5K | 116101 | 0.534642969 | 0 | YWHAE |
| 184 | 396316 | rs28365859 | chr17-1400484 | 0.165 | MYO1C | 90677 | 0.532800396 | 1 | YWHAE |
| 185 | 396316 | rs28365859 | chr17-1400484 | 0.165 | PITPNA-AS1 | 116371 | 0.519511352 | 2 | YWHAE |
| 186 | 398790 | rs9908102 | chr17-12993236 | 0.067 | ELAC2 | 24600 | 0.535925249 | 0 | ELAC2 |
| 187 | 399903 | rs959071 | chr17-19238913 | 0.382 | EPN2-AS1 | 67032 | 0.518669111 | 0 | EPN2 |
| 188 | 399903 | rs959071 | chr17-19238913 | 0.382 | B9D1 | 138964 | 0.507006818 | 1 | EPN2 |
| 189 | 399947 | rs12602286 | chr17-19333641 | 0.172 | MFAP4 | 53062 | 0.553732851 | 0 | EPN2-AS1 |
| 190 | 399947 | rs12602286 | chr17-19333641 | 0.172 | EPN2-AS1 | 27367 | 0.546348867 | 1 | EPN2-AS1 |
| 191 | 399947 | rs12602286 | chr17-19333641 | 0.172 | B9D1 | 44065 | 0.538736664 | 2 | EPN2-AS1 |
| 192 | 407176 | rs34888090 | chr17-57664675 | 0.088 | MSI2 | 10078 | 0.507750863 | 0 | CCDC182 |
| 193 | 407178 | rs12938078 | chr17-57665916 | 0.073 | MRPS23 | 183806 | 0.515961292 | 0 | CCDC182 |
| 194 | 407178 | rs12938078 | chr17-57665916 | 0.073 | MSI2 | 8781 | 0.513769988 | 1 | CCDC182 |
| 195 | 408133 | rs4291 | chr17-63476833 | 0.087 | CYB561 | 35590 | 0.547801473 | 0 | ACE |
| 196 | 408133 | rs4291 | chr17-63476833 | 0.087 | LIMD2 | 224243 | 0.523080934 | 1 | ACE |
| 197 | 408133 | rs4291 | chr17-63476833 | 0.087 | TACO1 | 124646 | 0.519911228 | 2 | ACE |
| 198 | 408136 | rs4295 | chr17-63478937 | 0.053 | CYB561 | 37984 | 0.543418646 | 0 | ACE |
| 199 | 408136 | rs4295 | chr17-63478937 | 0.053 | TACO1 | 122252 | 0.517891458 | 1 | ACE |

| | A | B | C | D | E | F | G | H | I |
|---|---|---|---|---|---|---|---|---|---|
| 200 | 408136 | rs4295 | chr17-63478937 | 0.053 | LIMD2 | 221849 | 0.516083783 | 2 | ACE |
| 201 | 422428 | rs4128242 | chr18-55080458 | 0.095 | RAB27B | 195621 | 0.511293109 | 0 | LINC01929 |
| 202 | 422429 | rs12969453 | chr18-55084477 | 0.121 | RAB27B | 199757 | 0.518613879 | 0 | LINC01929 |
| 203 | 422561 | rs74914300 | chr18-55343117 | 0.058 | TCF4-AS2 | 148988 | 0.526100813 | 0 | TCF4-AS1 |
| 204 | 422592 | rs75037433 | chr18-55390864 | 0.056 | TCF4-AS2 | 100863 | 0.532477931 | 0 | TCF4-AS1 |
| 205 | 422592 | rs75037433 | chr18-55390864 | 0.056 | TCF4-AS1 | 61221 | 0.510827016 | 1 | TCF4-AS1 |
| 206 | 422598 | rs78322266 | chr18-55396445 | 0.064 | TCF4-AS2 | 95420 | 0.532373459 | 0 | TCF4-AS1 |
| 207 | 422598 | rs78322266 | chr18-55396445 | 0.064 | TCF4-AS1 | 55778 | 0.513108253 | 1 | TCF4-AS1 |
| 208 | 426658 | rs509573 | chr18-79800879 | 0.107 | KCNG2 | 74619 | 0.53759408 | 0 | KCNG2 |
| 209 | 426658 | rs509573 | chr18-79800879 | 0.107 | SLC66A2 | 150392 | 0.519656695 | 1 | KCNG2 |
| 210 | 426658 | rs509573 | chr18-79800879 | 0.107 | CTDP1 | 64412 | 0.506286845 | 2 | KCNG2 |
| 211 | 426663 | rs4798923 | chr18-79869373 | 0.07 | KCNG2 | 6133 | 0.570059436 | 0 | KCNG2 |
| 212 | 426663 | rs4798923 | chr18-79869373 | 0.07 | HSBP1L1 | 100292 | 0.533798604 | 1 | KCNG2 |
| 213 | 426663 | rs4798923 | chr18-79869373 | 0.07 | SLC66A2 | 81906 | 0.528829274 | 2 | KCNG2 |
| 214 | 426664 | rs11660941 | chr18-79870600 | 0.07 | KCNG2 | 5051 | 0.578060168 | 0 | KCNG2 |
| 215 | 426664 | rs11660941 | chr18-79870600 | 0.07 | SLC66A2 | 80824 | 0.537551811 | 1 | KCNG2 |
| 216 | 426664 | rs11660941 | chr18-79870600 | 0.07 | HSBP1L1 | 99210 | 0.533140189 | 2 | KCNG2 |

| | A | B | C | D | E | F | G | H | I |
|---|---|---|---|---|---|---|---|---|---|
| 217 | 426665 | rs61090726 | chr18-79871219 | 0.063 | KCNG2 | 4367 | 0.575771059 | 0 | KCNG2 |
| 218 | 426665 | rs61090726 | chr18-79871219 | 0.063 | SLC66A2 | 80140 | 0.537748608 | 1 | KCNG2 |
| 219 | 426665 | rs61090726 | chr18-79871219 | 0.063 | HSBP1L1 | 98526 | 0.537262121 | 2 | KCNG2 |
| 220 | 426666 | rs72980085 | chr18-79871679 | 0.08 | KCNG2 | 3769 | 0.577368456 | 0 | KCNG2 |
| 221 | 426666 | rs72980085 | chr18-79871679 | 0.08 | SLC66A2 | 79542 | 0.545208017 | 1 | KCNG2 |
| 222 | 426666 | rs72980085 | chr18-79871679 | 0.08 | HSBP1L1 | 97928 | 0.533410957 | 2 | KCNG2 |
| 223 | 426667 | rs8091497 | chr18-79872565 | 0.09 | KCNG2 | 2873 | 0.560140138 | 0 | KCNG2 |
| 224 | 426667 | rs8091497 | chr18-79872565 | 0.09 | SLC66A2 | 78646 | 0.538529109 | 1 | KCNG2 |
| 225 | 426667 | rs8091497 | chr18-79872565 | 0.09 | HSBP1L1 | 97032 | 0.518167152 | 2 | KCNG2 |
| 226 | 430431 | rs322124 | chr19-11288696 | 0.239 | DOCK6 | 34172 | 0.554087633 | 0 | TSPAN16 |
| 227 | 430431 | rs322124 | chr19-11288696 | 0.239 | TMEM205 | 57542 | 0.550831696 | 1 | TSPAN16 |
| 228 | 430431 | rs322124 | chr19-11288696 | 0.239 | CCDC159 | 63242 | 0.548895192 | 2 | TSPAN16 |
| 229 | 430433 | rs322129 | chr19-11290429 | 0.223 | DOCK6 | 36117 | 0.543044544 | 0 | TSPAN16 |
| 230 | 430433 | rs322129 | chr19-11290429 | 0.223 | CCDC159 | 61297 | 0.541698906 | 1 | TSPAN16 |
| 231 | 430433 | rs322129 | chr19-11290429 | 0.223 | TMEM205 | 55597 | 0.535393988 | 2 | TSPAN16 |
| 232 | 430560 | rs72986630 | chr19-11738921 | 0.99 | ZNF823 | 247 | 0.559184299 | 0 | ZNF823 |
| 233 | 430560 | rs72986630 | chr19-11738921 | 0.99 | ZNF441 | 27744 | 0.552197366 | 1 | ZNF823 |

| | A | B | C | D | E | F | G | H | I |
|---|---|---|---|---|---|---|---|---|---|
| 234 | 430560 | rs72986630 | chr19-11738921 | 0.99 | ZNF491 | 59332 | 0.534652146 | 2 | ZNF823 |
| 235 | 437940 | rs2304204 | chr19-49665763 | 0.09 | SCAF1 | 20729 | 0.558311257 | 0 | IRF3 |
| 236 | 437940 | rs2304204 | chr19-49665763 | 0.09 | IRF3 | 102 | 0.548925285 | 1 | IRF3 |
| 237 | 437940 | rs2304204 | chr19-49665763 | 0.09 | CPT1C | 46841 | 0.547245293 | 2 | IRF3 |
| 238 | 439190 | rs758749 | chr19-56678350 | 0.346 | ZNF71 | 82771 | 0.547101722 | 0 | ZNF835 |
| 239 | 439190 | rs758749 | chr19-56678350 | 0.346 | MIMT1 | 162643 | 0.540521334 | 1 | ZNF835 |
| 240 | 439190 | rs758749 | chr19-56678350 | 0.346 | ZNF835 | 6323 | 0.538772949 | 2 | ZNF835 |

| | A | B | C | D | E | F |
|---|---|---|---|---|---|---|
| 1 | **Table S2** | | | | | |
| 2 | gene_from_gridnet | tf | log10VariantEffect_from_FIMO | position | rsid_from_PMID_35396580 | effect_on_binding_affinity |
| 3 | TBL1XR1-AS1 | KLF15 | -4.07849 | chr3-176767108 | rs9825834 | TF loss |
| 4 | TBL1XR1-AS1 | SP5 | -3.89992 | chr3-176767108 | rs9825834 | TF loss |
| 5 | TBL1XR1-AS1 | MAZ | -3.69357 | chr3-176767108 | rs9825834 | TF loss |
| 6 | TBL1XR1-AS1 | MAZ | -3.31091 | chr3-176767108 | rs9825834 | TF loss |
| 7 | TBL1XR1-AS1 | SP1 | -2.72236 | chr3-176767108 | rs9825834 | TF loss |
| 8 | MAP3K11, PCNX3, RNASEH2C | KLF6 | 2.640419 | chr11-65383755 | rs58950470 | TF gain |
| 9 | TBL1XR1-AS1 | ZN263 | 2.609962 | chr3-176767108 | rs9825834 | TF gain |
| 10 | MAP3K11, PCNX3, RNASEH2C | PLAG1 | -2.5616 | chr11-65383755 | rs58950470 | TF loss |
| 11 | TBL1XR1-AS1 | KLF15 | -2.55654 | chr3-176767108 | rs9825834 | TF loss |
| 12 | SLC12A6, LPCAT4, NOP10 | KLF6 | -2.52533 | chr15-34659517 | rs117799466 | TF loss |
| 13 | TBL1XR1-AS1 | VEZF1 | -2.51755 | chr3-176767108 | rs9825834 | TF loss |
| 14 | TBL1XR1-AS1 | KLF1 | -2.40514 | chr3-176767108 | rs9825834 | TF loss |
| 15 | TBL1XR1-AS1 | SP1 | -2.40302 | chr3-176767108 | rs9825834 | TF loss |

| | A | B | C | D | E | F |
|---|---|---|---|---|---|---|
| 16 | NAB2, STAT6, NXPH4 | PURA | -2.39794 | chr12-57487814 | rs324017 | TF loss |
| 17 | TBL1XR1-AS1 | KLF5 | -2.38352 | chr3-176767108 | rs9825834 | TF loss |
| 18 | SLC12A6, LPCAT4, NOP10 | SP1 | 2.373195 | chr15-34659517 | rs117799466 | TF gain |
| 19 | TBL1XR1-AS1 | SP3 | -2.35482 | chr3-176767108 | rs9825834 | TF loss |
| 20 | NAB2, STAT6, NXPH4 | PURA | -2.33488 | chr12-57487814 | rs324017 | TF loss |
| 21 | SLC12A6, LPCAT4, NOP10 | ZN281 | 2.232606 | chr15-34659517 | rs117799466 | TF gain |
| 22 | MAP3K11, PCNX3, RNASEH2C | WT1 | -2.2213 | chr11-65383755 | rs58950470 | TF loss |
| 23 | MAP3K11, PCNX3, RNASEH2C | SP1 | -2.21927 | chr11-65383755 | rs58950470 | TF loss |
| 24 | MAP3K11, PCNX3, RNASEH2C | SP3 | -2.2165 | chr11-65383755 | rs58950470 | TF loss |
| 25 | TBL1XR1-AS1 | ZNF281 | -2.19735 | chr3-176767108 | rs9825834 | TF loss |
| 26 | TBL1XR1-AS1 | KLF16 | -2.1916 | chr3-176767108 | rs9825834 | TF loss |
| 27 | SLC12A6, LPCAT4, NOP10 | MAZ | 2.183573 | chr15-34659517 | rs117799466 | TF gain |
| 28 | TBL1XR1-AS1 | PATZ1 | -2.10984 | chr3-176767108 | rs9825834 | TF loss |
| 29 | ARNILA, DTNB-AS1 | KLF6 | -2.10134 | chr2-25438963 | rs527882868 | TF loss |
| 30 | SLC12A6, LPCAT4, NOP10 | SP2 | 2.095089 | chr15-34659517 | rs117799466 | TF gain |

| | A | B | C | D | E | F |
|---|---|---|---|---|---|---|
| 31 | MAP3K11, PCNX3, RNASEH2C | PATZ1 | -2.07479 | chr11-65383755 | rs58950470 | TF loss |
| 32 | ICA1 | ZNF440 | 2.074119 | chr7-8549187 | rs11972718 | TF gain |
| 33 | SLC12A6, LPCAT4, NOP10 | HKR1 | 2.051694 | chr15-34659517 | rs117799466 | TF gain |
| 34 | TBL1XR1-AS1 | SP2 | -2.01878 | chr3-176767108 | rs9825834 | TF loss |
| 35 | EPHX2 | KLF6 | -2.00421 | chr8-27328354 | rs2472554 | TF loss |
| 36 | SLC12A6, LPCAT4, NOP10 | SP4 | 2.002719 | chr15-34659517 | rs117799466 | TF gain |
| 37 | TBL1XR1-AS1 | KLF1 | -1.98038 | chr3-176767108 | rs9825834 | TF loss |
| 38 | TBL1XR1-AS1 | KLF15 | 1.978058 | chr3-176767108 | rs9825834 | TF gain |
| 39 | TBL1XR1-AS1 | KLF6 | -1.9778 | chr3-176767108 | rs9825834 | TF loss |
| 40 | MAP3K11, PCNX3, RNASEH2C | ZNF37A | -1.97126 | chr11-65383755 | rs58950470 | TF loss |
| 41 | MAP3K11, PCNX3, RNASEH2C | SP3 | -1.96586 | chr11-65383755 | rs58950470 | TF loss |
| 42 | SLC12A6, LPCAT4, NOP10 | OSR2 | -1.95823 | chr15-34659517 | rs117799466 | TF loss |
| 43 | SLC12A6, LPCAT4, NOP10 | ZN281 | 1.896833 | chr15-34659517 | rs117799466 | TF gain |
| 44 | SLC12A6, LPCAT4, NOP10 | SP4 | 1.886301 | chr15-34659517 | rs117799466 | TF gain |
| 45 | MAP3K11, PCNX3, RNASEH2C | MAZ | -1.87932 | chr11-65383755 | rs58950470 | TF loss |

| | A | B | C | D | E | F |
|---|---|---|---|---|---|---|
| 46 | SLC12A6, LPCAT4, NOP10 | SP1 | 1.875827 | chr15-34659517 | rs117799466 | TF gain |
| 47 | TBL1XR1-AS1 | KLF12 | -1.86923 | chr3-176767108 | rs9825834 | TF loss |
| 48 | SLC12A6, LPCAT4, NOP10 | ZSC22 | 1.867134 | chr15-34659517 | rs117799466 | TF gain |
| 49 | MAP3K11, PCNX3, RNASEH2C | ZNF707 | 1.848676 | chr11-65383755 | rs58950470 | TF gain |
| 50 | EMX1 | VEZF1 | -1.84253 | chr2-73168593 | rs6722190 | TF loss |
| 51 | TBL1XR1-AS1 | MAZ | 1.84018 | chr3-176767108 | rs9825834 | TF gain |
| 52 | SLC12A6, LPCAT4, NOP10 | WT1 | 1.830145 | chr15-34659517 | rs117799466 | TF gain |
| 53 | MAP3K11, PCNX3, RNASEH2C | ZNF148 | -1.82925 | chr11-65383755 | rs58950470 | TF loss |
| 54 | MAP3K11, PCNX3, RNASEH2C | SP1 | -1.82729 | chr11-65383755 | rs58950470 | TF loss |
| 55 | SLC12A6, LPCAT4, NOP10 | VEZF1 | 1.826595 | chr15-34659517 | rs117799466 | TF gain |
| 56 | MAP3K11, PCNX3, RNASEH2C | SP3 | -1.81299 | chr11-65383755 | rs58950470 | TF loss |
| 57 | MAP3K11, PCNX3, RNASEH2C | TBX15 | -1.80695 | chr11-65383755 | rs58950470 | TF loss |
| 58 | SLC12A6, LPCAT4, NOP10 | ZNF148 | 1.793663 | chr15-34659517 | rs117799466 | TF gain |
| 59 | SLC12A6, LPCAT4, NOP10 | ZBT17 | 1.785063 | chr15-34659517 | rs117799466 | TF gain |
| 60 | SLC12A6, LPCAT4, NOP10 | PATZ1 | 1.774062 | chr15-34659517 | rs117799466 | TF gain |

| | A | B | C | D | E | F |
|---|---|---|---|---|---|---|
| 61 | NAB2, STAT6, NXPH4 | ZNF780A | -1.76737 | chr12-57487814 | rs324017 | TF loss |
| 62 | NAB2, STAT6, NXPH4 | ZN335 | -1.75868 | chr12-57487814 | rs324017 | TF loss |
| 63 | MAP3K11, PCNX3, RNASEH2C | KLF3 | 1.752558 | chr11-65383755 | rs58950470 | TF gain |
| 64 | SLC12A6, LPCAT4, NOP10 | PRDM9 | 1.727846 | chr15-34659517 | rs117799466 | TF gain |
| 65 | MAP3K11, PCNX3, RNASEH2C | MAZ | -1.7275 | chr11-65383755 | rs58950470 | TF loss |
| 66 | TBL1XR1-AS1 | WT1 | -1.72163 | chr3-176767108 | rs9825834 | TF loss |
| 67 | TBL1XR1-AS1 | ZN467 | -1.72141 | chr3-176767108 | rs9825834 | TF loss |
| 68 | TBL1XR1-AS1 | ZNF281 | -1.72127 | chr3-176767108 | rs9825834 | TF loss |
| 69 | SLC12A6, LPCAT4, NOP10 | SP4 | -1.71551 | chr15-34659517 | rs117799466 | TF loss |
| 70 | KLF10 | SP4 | -1.71018 | chr8-103669991 | rs4734654 | TF loss |
| 71 | DLX2, DLX1 | SP2 | 1.708357 | chr2-172968775 | rs62183859 | TF gain |
| 72 | MAP3K11, PCNX3, RNASEH2C | SP1 | -1.70466 | chr11-65383755 | rs58950470 | TF loss |
| 73 | MAP3K11, PCNX3, RNASEH2C | SP2 | -1.69734 | chr11-65383755 | rs58950470 | TF loss |
| 74 | SLC12A6, LPCAT4, NOP10 | ZNF281 | 1.697203 | chr15-34659517 | rs117799466 | TF gain |
| 75 | C12orf43, P2RX7 | ZNF610 | -1.68518 | chr12-121341165 | rs7486605 | TF loss |

| | A | B | C | D | E | F |
|---|---|---|---|---|---|---|
| 76 | SFXN2, WBP1L, C10orf95-AS1 | ZN713 | 1.670559 | chr10-104420018 | rs10883761 | TF gain |
| 77 | ARNILA, DTNB-AS1 | KLF1 | 1.66549 | chr2-25438963 | rs527882868 | TF gain |
| 78 | MAP3K11, PCNX3, RNASEH2C | ZN148 | -1.65392 | chr11-65383755 | rs58950470 | TF loss |
| 79 | SLC12A6, LPCAT4, NOP10 | SP3 | 1.651109 | chr15-34659517 | rs117799466 | TF gain |
| 80 | SLC12A6, LPCAT4, NOP10 | SP2 | 1.65042 | chr15-34659517 | rs117799466 | TF gain |
| 81 | MAP3K11, PCNX3, RNASEH2C | MAZ | -1.6503 | chr11-65383755 | rs58950470 | TF loss |
| 82 | MAP3K11, PCNX3, RNASEH2C | ZN263 | -1.64724 | chr11-65383755 | rs58950470 | TF loss |
| 83 | MFAP4, EPN2-AS1, B9D1 | ZNF267 | -1.63259 | chr17-19236954 | rs12602286 | TF loss |
| 84 | MAP3K11, PCNX3, RNASEH2C | MAZ | -1.62598 | chr11-65383755 | rs58950470 | TF loss |
| 85 | EPHX2 | KLF16 | 1.619689 | chr8-27328354 | rs2472554 | TF gain |
| 86 | DLX2, DLX1 | ZFX | -1.61621 | chr2-172974789 | rs62184960 | TF loss |
| 87 | SLC12A6, LPCAT4, NOP10 | SP3 | 1.606717 | chr15-34659517 | rs117799466 | TF gain |
| 88 | SFXN2, WBP1L, C10orf95-AS1 | ZNF823 | 1.606381 | chr10-104420018 | rs10883761 | TF gain |
| 89 | MAP3K11, PCNX3, RNASEH2C | ZNF460 | -1.60594 | chr11-65383755 | rs58950470 | TF loss |
| 90 | TBL1XR1-AS1 | ZNF148 | -1.60128 | chr3-176767108 | rs9825834 | TF loss |

| | A | B | C | D | E | F |
|---|---|---|---|---|---|---|
| 91 | KCNG2, SLC66A2, HSBP1L1 | SP1 | -1.60085 | chr18-77632565 | rs8091497 | TF loss |
| 92 | SLC12A6, LPCAT4, NOP10 | VEZF1 | 1.59936 | chr15-34659517 | rs117799466 | TF gain |
| 93 | MAP3K11, PCNX3, RNASEH2C | KLF3 | -1.59146 | chr11-65383755 | rs58950470 | TF loss |
| 94 | TBL1XR1-AS1 | KLF6 | 1.589185 | chr3-176767108 | rs9825834 | TF gain |
| 95 | ZNF71, MIMT1, ZNF835 | ZNF610 | -1.58893 | chr19-57189718 | rs758749 | TF loss |
| 96 | TBL1XR1-AS1 | VEZF1 | 1.588413 | chr3-176767108 | rs9825834 | TF gain |
| 97 | TBL1XR1-AS1 | ZN281 | -1.58827 | chr3-176767108 | rs9825834 | TF loss |
| 98 | TBL1XR1-AS1 | ZNF398 | 1.585789 | chr3-176767108 | rs9825834 | TF gain |
| 99 | MAP3K11, PCNX3, RNASEH2C | ZNF467 | -1.58246 | chr11-65383755 | rs58950470 | TF loss |
| 100 | SLC12A6, LPCAT4, NOP10 | EGR2 | 1.567812 | chr15-34659517 | rs117799466 | TF gain |
| 101 | SLC12A6, LPCAT4, NOP10 | ZNF281 | 1.548375 | chr15-34659517 | rs117799466 | TF gain |
| 102 | SLC12A6, LPCAT4, NOP10 | ZNF460 | 1.539133 | chr15-34659517 | rs117799466 | TF gain |
| 103 | DLX2, DLX1 | SP3 | 1.53844 | chr2-172968775 | rs62183859 | TF gain |
| 104 | TBL1XR1-AS1 | ZN148 | -1.53494 | chr3-176767108 | rs9825834 | TF loss |
| 105 | SLC12A6, LPCAT4, NOP10 | EGR1 | 1.527402 | chr15-34659517 | rs117799466 | TF gain |

| | A | B | C | D | E | F |
|---|---|---|---|---|---|---|
| 106 | DLX2, DLX1 | KLF15 | -1.51964 | chr2-172968775 | rs62183859 | TF loss |
| 107 | KLF10 | TBX1 | 1.511588 | chr8-103669991 | rs4734654 | TF gain |
| 108 | MAP3K11, PCNX3, RNASEH2C | KLF15 | -1.50922 | chr11-65383755 | rs58950470 | TF loss |
| 109 | NAB2, STAT6, NXPH4 | ZN335 | -1.50775 | chr12-57487814 | rs324017 | TF loss |
| 110 | TBL1XR1-AS1 | MAZ | 1.505464 | chr3-176767108 | rs9825834 | TF gain |
| 111 | DLX2, DLX1 | KLF16 | 1.500165 | chr2-172968775 | rs62183859 | TF gain |
| 112 | ARNILA, DTNB-AS1 | SP1 | 1.499813 | chr2-25438963 | rs527882868 | TF gain |
| 113 | TBL1XR1-AS1 | MAZ | -1.49971 | chr3-176767108 | rs9825834 | TF loss |
| 114 | PLCH2, PEX10, MORN1 | SP3 | 1.499231 | chr1-2374758 | rs4648844 | TF gain |
| 115 | MAP3K11, PCNX3, RNASEH2C | TBX15 | -1.49917 | chr11-65383755 | rs58950470 | TF loss |
| 116 | KLF10 | KLF15 | -1.49714 | chr8-103669991 | rs4734654 | TF loss |
| 117 | TBL1XR1-AS1 | MAZ | -1.49334 | chr3-176767108 | rs9825834 | TF loss |
| 118 | KCNG2, SLC66A2, HSBP1L1 | ZN770 | -1.48734 | chr18-77631679 | rs72980085 | TF loss |
| 119 | SLC12A6, LPCAT4, NOP10 | Wt1 | 1.486498 | chr15-34659517 | rs117799466 | TF gain |
| 120 | SLC12A6, LPCAT4, NOP10 | SP2 | -1.48578 | chr15-34659517 | rs117799466 | TF loss |

| | A | B | C | D | E | F |
|---|---|---|---|---|---|---|
| 121 | SLC12A6, LPCAT4, NOP10 | WT1 | 1.483945 | chr15-34659517 | rs117799466 | TF gain |
| 122 | TBL1XR1-AS1 | ZNF468 | -1.4829 | chr3-176767108 | rs9825834 | TF loss |
| 123 | ZNF202 | ZNF341 | -1.47893 | chr11-123395987 | rs7927437 | TF loss |
| 124 | PLCH2, PEX10, MORN1 | VEZF1 | -1.47815 | chr1-2374758 | rs4648844 | TF loss |
| 125 | TBL1XR1-AS1 | ZNF148 | 1.47814 | chr3-176767108 | rs9825834 | TF gain |
| 126 | MAP3K11, PCNX3, RNASEH2C | E2F1 | -1.47799 | chr11-65383755 | rs58950470 | TF loss |
| 127 | EPHX2 | ZNF132 | 1.475882 | chr8-27328354 | rs2472554 | TF gain |
| 128 | SLC12A6, LPCAT4, NOP10 | TBX15 | 1.475587 | chr15-34659517 | rs117799466 | TF gain |
| 129 | MYT1L-AS1 | TBX15 | 1.473191 | chr2-2330118 | rs6708950 | TF gain |
| 130 | PLCH2, PEX10, MORN1 | SP2 | 1.472712 | chr1-2374758 | rs4648844 | TF gain |
| 131 | NAB2, STAT6, NXPH4 | ZN770 | -1.4719 | chr12-57487814 | rs324017 | TF loss |
| 132 | MAP3K11, PCNX3, RNASEH2C | KLF12 | -1.46699 | chr11-65383755 | rs58950470 | TF loss |
| 133 | MAP3K11, PCNX3, RNASEH2C | ZNF460 | -1.46152 | chr11-65383755 | rs58950470 | TF loss |
| 134 | SLC12A6, LPCAT4, NOP10 | MAZ | 1.461301 | chr15-34659517 | rs117799466 | TF gain |
| 135 | TBL1XR1-AS1 | TBX15 | -1.45976 | chr3-176767108 | rs9825834 | TF loss |
| 136 | IGSF9B, THYN1 | ZNF282 | 1.459585 | chr11-133905334 | rs687187 | TF gain |

| | A | B | C | D | E | F |
|---|---|---|---|---|---|---|
| 137 | SLC12A6, LPCAT4, NOP10 | EGR2 | 1.459077 | chr15-34659517 | rs117799466 | TF gain |
| 138 | SLC12A6, LPCAT4, NOP10 | ZNF774 | 1.457779 | chr15-34659517 | rs117799466 | TF gain |
| 139 | DLX2, DLX1 | ZNF180 | 1.455626 | chr2-172968775 | rs62183859 | TF gain |
| 140 | SLC12A6, LPCAT4, NOP10 | SP1 | 1.450544 | chr15-34659517 | rs117799466 | TF gain |
| 141 | ARNILA, DTNB-AS1 | ZNF534 | 1.449192 | chr2-25438963 | rs527882868 | TF gain |
| 142 | MAP3K11, PCNX3, RNASEH2C | ZNF341 | -1.44909 | chr11-65383755 | rs58950470 | TF loss |
| 143 | TBL1XR1-AS1 | ZN263 | -1.44768 | chr3-176767108 | rs9825834 | TF loss |
| 144 | ARNILA, DTNB-AS1 | KLF3 | -1.44569 | chr2-25438963 | rs527882868 | TF loss |
| 145 | ASIC1, COX14, NCKAP5L | ZNF320 | 1.44364 | chr12-50464470 | rs706790 | TF gain |
| 146 | MAP3K11, PCNX3, RNASEH2C | ZNF180 | -1.44305 | chr11-65383755 | rs58950470 | TF loss |
| 147 | MAP3K11, PCNX3, RNASEH2C | E2F1 | -1.44267 | chr11-65383755 | rs58950470 | TF loss |
| 148 | KLF10 | VEZF1 | -1.4415 | chr8-103669991 | rs4734654 | TF loss |
| 149 | MAP3K11, PCNX3, RNASEH2C | ZNF468 | -1.43806 | chr11-65383755 | rs58950470 | TF loss |
| 150 | SLC12A6, LPCAT4, NOP10 | EGR2 | 1.436503 | chr15-34659517 | rs117799466 | TF gain |
| 151 | SLC12A6, LPCAT4, NOP10 | ZNF281 | 1.435423 | chr15-34659517 | rs117799466 | TF gain |

| | A | B | C | D | E | F |
|---|---|---|---|---|---|---|
| 152 | MRPS23, MSI2 | ZNF580 | -1.43537 | chr17-55743277 | rs12938078 | TF loss |
| 153 | SLC12A6, LPCAT4, NOP10 | EGR2 | 1.425051 | chr15-34659517 | rs117799466 | TF gain |
| 154 | TBL1XR1-AS1 | ZNF467 | -1.41899 | chr3-176767108 | rs9825834 | TF loss |
| 155 | NAB2, STAT6, NXPH4 | PRDM9 | -1.41452 | chr12-57487814 | rs324017 | TF loss |
| 156 | EMX1 | MAZ | -1.41173 | chr2-73168593 | rs6722190 | TF loss |
| 157 | TBL1XR1-AS1 | ZNF467 | 1.40981 | chr3-176767108 | rs9825834 | TF gain |
| 158 | MAP3K11, PCNX3, RNASEH2C | ZN148 | -1.40902 | chr11-65383755 | rs58950470 | TF loss |
| 159 | SLC12A6, LPCAT4, NOP10 | KLF15 | 1.406484 | chr15-34659517 | rs117799466 | TF gain |
| 160 | TBL1XR1-AS1 | ZN467 | -1.39717 | chr3-176767108 | rs9825834 | TF loss |
| 161 | TBL1XR1-AS1 | SP4 | -1.39332 | chr3-176767108 | rs9825834 | TF loss |
| 162 | KLF10 | ZBT17 | -1.39222 | chr8-103669991 | rs4734654 | TF loss |
| 163 | ARNILA, DTNB-AS1 | SP1 | 1.391814 | chr2-25438963 | rs527882868 | TF gain |
| 164 | PLCH2, PEX10, MORN1 | TBX1 | 1.391458 | chr1-2374758 | rs4648844 | TF gain |
| 165 | TBL1XR1-AS1 | ZN467 | 1.38489 | chr3-176767108 | rs9825834 | TF gain |
| 166 | SLC12A6, LPCAT4, NOP10 | SP2 | 1.384542 | chr15-34659517 | rs117799466 | TF gain |

| | A | B | C | D | E | F |
|---|---|---|---|---|---|---|
| 167 | TAF12 | ZNF496 | -1.38102 | chr1-29150658 | rs12092160 | TF loss |
| 168 | TBL1XR1-AS1 | SP5 | 1.380485 | chr3-176767108 | rs9825834 | TF gain |
| 169 | MAP3K11, PCNX3, RNASEH2C | PLAG1 | -1.37827 | chr11-65383755 | rs58950470 | TF loss |
| 170 | NAB2, STAT6, NXPH4 | HKR1 | -1.37727 | chr12-57487814 | rs324017 | TF loss |
| 171 | KCNG2, SLC66A2, CTDP1 | ZN341 | -1.37634 | chr18-77560879 | rs509573 | TF loss |
| 172 | MAP3K11, PCNX3, RNASEH2C | PATZ1 | -1.37358 | chr11-65383755 | rs58950470 | TF loss |
| 173 | SLC12A6, LPCAT4, NOP10 | SP2 | 1.370303 | chr15-34659517 | rs117799466 | TF gain |
| 174 | PLCH2, PEX10, MORN1 | PURA | 1.367851 | chr1-2374758 | rs4648844 | TF gain |
| 175 | TBL1XR1-AS1 | ZNF444 | -1.3637 | chr3-176767108 | rs9825834 | TF loss |
| 176 | MAP3K11, PCNX3, RNASEH2C | CTCFL | -1.36228 | chr11-65383755 | rs58950470 | TF loss |
| 177 | MRPS23, MSI2 | KLF16 | -1.3614 | chr17-55743277 | rs12938078 | TF loss |
| 178 | TBL1XR1-AS1 | KLF1 | 1.361376 | chr3-176767108 | rs9825834 | TF gain |
| 179 | PSD2 | ZNF571 | -1.36103 | chr5-139028810 | rs10447226 | TF loss |
| 180 | NAB2, STAT6, NXPH4 | KLF3 | -1.36057 | chr12-57487814 | rs324017 | TF loss |
| 181 | SLC12A6, LPCAT4, NOP10 | SP3 | 1.359506 | chr15-34659517 | rs117799466 | TF gain |

| | A | B | C | D | E | F |
|---|---|---|---|---|---|---|
| 182 | MAP3K11, PCNX3, RNASEH2C | ZN341 | -1.35485 | chr11-65383755 | rs58950470 | TF loss |
| 183 | EPHX2 | MAZ | 1.354589 | chr8-27328354 | rs2472554 | TF gain |
| 184 | DLX2, DLX1 | SP1 | 1.354506 | chr2-172968775 | rs62183859 | TF gain |
| 185 | SLC12A6, LPCAT4, NOP10 | KLF16 | 1.354432 | chr15-34659517 | rs117799466 | TF gain |
| 186 | SLC12A6, LPCAT4, NOP10 | KLF15 | 1.353072 | chr15-34659517 | rs117799466 | TF gain |
| 187 | PLCH2, PEX10, MORN1 | SPIC | 1.346787 | chr1-2374758 | rs4648844 | TF gain |
| 188 | EPHX2 | ZNF398 | 1.345411 | chr8-27328354 | rs2472554 | TF gain |
| 189 | SLC12A6, LPCAT4, NOP10 | SP2 | 1.34034 | chr15-34659517 | rs117799466 | TF gain |
| 190 | SLC12A6, LPCAT4, NOP10 | SP4 | 1.339904 | chr15-34659517 | rs117799466 | TF gain |
| 191 | PLCH2, PEX10, MORN1 | ZNF28 | 1.337427 | chr1-2374758 | rs4648844 | TF gain |
| 192 | TBL1XR1-AS1 | ZN148 | -1.3248 | chr3-176767108 | rs9825834 | TF loss |
| 193 | SLC12A6, LPCAT4, NOP10 | SP2 | -1.3248 | chr15-34659517 | rs117799466 | TF loss |
| 194 | SLC12A6, LPCAT4, NOP10 | TBX15 | 1.323596 | chr15-34659517 | rs117799466 | TF gain |
| 195 | NAB2, STAT6, NXPH4 | SP3 | -1.32201 | chr12-57487814 | rs324017 | TF loss |
| 196 | MAP3K11, PCNX3, RNASEH2C | ZN467 | -1.32118 | chr11-65383755 | rs58950470 | TF loss |
| 197 | DLX2, DLX1 | Zfx | -1.32086 | chr2-172974789 | rs62184960 | TF loss |

| | A | B | C | D | E | F |
|---|---|---|---|---|---|---|
| 198 | PSD2 | SP2 | -1.32057 | chr5-139028810 | rs10447226 | TF loss |
| 199 | DLX2, DLX1 | SP3 | 1.320193 | chr2-172968775 | rs62183859 | TF gain |
| 200 | NAB2, STAT6, NXPH4 | VEZF1 | -1.31778 | chr12-57487814 | rs324017 | TF loss |
| 201 | EPHX2 | ZBT17 | 1.314125 | chr8-27328354 | rs2472554 | TF gain |
| 202 | MAP3K11, PCNX3, RNASEH2C | MAZ | -1.3129 | chr11-65383755 | rs58950470 | TF loss |
| 203 | MAP3K11, PCNX3, RNASEH2C | ZN467 | -1.30883 | chr11-65383755 | rs58950470 | TF loss |
| 204 | KLF10 | PRDM9 | 1.308454 | chr8-103669991 | rs4734654 | TF gain |
| 205 | SLC12A6, LPCAT4, NOP10 | HKR1 | 1.307877 | chr15-34659517 | rs117799466 | TF gain |
| 206 | DLX2, DLX1, METAP1D | ZNF341 | -1.30621 | chr2-172955433 | rs62183854 | TF loss |
| 207 | TBL1XR1-AS1 | VEZF1 | 1.30459 | chr3-176767108 | rs9825834 | TF gain |
| 208 | ZEB2-AS1, GTDC1 | ZNF823 | -1.3042 | chr2-145141541 | rs12991836 | TF loss |
| 209 | SLC12A6, LPCAT4, NOP10 | KLF15 | 1.30312 | chr15-34659517 | rs117799466 | TF gain |
